# The thermodynamic state of aerobic glycolytic flux plays a critical role in stabilizing aerobic glycolytic flux of cancer cells

**DOI:** 10.1101/2022.07.07.499167

**Authors:** Chengmeng Jin, Wei Hu, Yuqi Wang, Hao Wu, Siying Zeng, Minfeng Ying, Xun Hu

## Abstract

Inhibition of aerobic glycolysis (AG) is a strategy for cancer treatment, but its clinical translation has not been successful, suggesting that the mechanism underlying the inhibition of AG is not completely understood. We introduce thermodynamic state of AG (TSAG) into AG flux control, which refers the actual change of Gibbs free energy (ΔG) of reactions along the pathway. By using PKM2 knockdown and knockout models, we found that TSAG is resistant to PKM2 perturbation and it can effectively counteract the inhibitory effect of PKM2 perturbation on AG flux. Moreover, we deciphered the interrelationship between TSAG, AG rate, intermediate concentrations, and PK activity, revealed that TSAG is a system property of AG independent of cancer cells’ pathophysiological features and it plays important roles in stabilizing AG flux responding to perturbation. We propose that both inhibition of glycolytic enzyme and disruption of TSAG are required for effective AG inhibition.

## Introduction

AG or Warburg effect is a prominent metabolic feature of rapidly proliferating cancer cells, characterized by high glycolysis rates even under ample oxygen(Warburg, 1956). The high rate of glycolysis has 3 major metabolic purposes to support rapid growth of cancer cells(Vander Heiden, Cantley, & Thompson, 2009). First, the fast degradation of glucose transfers the energy to produce ATP. Second, the intermediates in the glycolysis flux shuttle to metabolic branches subsidiary to glycolysis for biosynthesis, for examples, G6P is used to generate NADPH and ribose-5-phospate through pentose phosphate pathway, DHAP is used to generate glycerol 3-phosphate, 3-PG is used to generate serine via serine synthesis pathway, and among others. Third, NADPH generated from oxidative branch of pentose phosphate pathway is the major reducing power not only for supporting biosynthesis but also for maintaining redox homeostasis.

Because of its importance in cancer cell metabolism, targeting AG has become a strategy for cancer treatment (Hanahan, 2022; Hanahan & Weinberg, 2011; Ward & Thompson, 2012). One approach to target cancer AG is to inhibit glycolytic enzymes. Numerous small molecule inhibitors for glycolytic enzymes were discovered and developed (X. S. Chen, Li, Guan, Yang, & Cheng, 2016; Kozal, Jozwiak, & Krzeslak, 2021; Tyagi, Mandal, & Roy, 2021). Nevertheless, the clinical translation by targeting glycolytic enzymes has not been encouraging(Kozal et al., 2021).

We may re-reconsider a basic question regarding the interrelationship between glycolytic enzymes and AG. There are mixed reports regarding roles of glycolytic enzymes in AG flux control. Many demonstrated that AG is sensitive to perturbation of glycolytic enzymes such as PKM2 (Anastasiou et al., 2012; Chaneton et al., 2012; Christofk, Vander Heiden, Harris, et al., 2008), PGK1(Hu et al., 2017; Y. Zhang et al., 2018), GAPDH (Liberti et al., 2017; Shestov et al., 2014), among others, while some reports that perturbation of glycolytic enzymes (PKM2, PGK1, GAPDH)(Jin, Zhu, Wu, Wang, & Hu, 2020; Tanner et al., 2018; Xie, Dai, & Hu, 2016; Zhu, Jin, Pan, & Hu, 2021) exerts no significant effects on AG. Then what causes this confliction?

We noted that previous researches mainly focus on enzyme kinetic part in AG but barely consider the thermodynamic part of AG. In theory, AG flux control is a question of kinetics and thermodynamics hence there should be an interaction between kinetics and thermodynamics. We recently reported that the thermodynamic factor indeed plays a role in stabilizing the actual activity of PGK and GAPDH in AG flux(Jin et al., 2020; Zhu et al., 2021): when PGK1 or GAPDH is inhibited by gene knockdown or inhibitor, the thermodynamic factor counteracts the inhibition, resulting in a stable rate through PGK1 or GAPDH, hence the overall AG rate is not disturbed. On this basis, we propose that thermodynamic part of AG is a long-overlooked factor and it plays an important role in stabilizing AG flux.

In this study, we found that TSAG is critical in AG flux control. Because PKM2 is classically considered as a rate-limiting enzyme of AG, we used PKM2 knockdown and knockout model to investigate the interrelationship between PKM2, AG flux, and TSAG. We found that TSAG can effectively counteract the effect of PKM2 knockdown or knockout on AG flux and we deciphered the underlying mechanism. This finding, in line with our previous reports (Jin et al., 2020; Zhu et al., 2021), provides a new thought for treating cancer by targeting AG, i.e., inhibiting AG should also take TSAG into consideration besides inhibiting glycolytic enzymes.

## Results

### Different cancer cells share a nearly identical TSAG

TSAG in cancer cells is defined by the concentrations of the intermediates, the reaction quotients (Qs) and the actual changes of the Gibbs free energy (ΔG) of the reactions along the glycolytic flux in cancer cells.

We measured glycolytic intermediates in different cancer cell lines from different origins (Human lung cancer cell line A549, cervical cancer cell line HeLa, gastric cancer cell line MGC80-3, liver cancer cell line SK-HEP-1, lung cancer cell line H1299 and colon cancer cell line RKO). The concentrations of the glycolytic intermediates varied between different cell lines (Figure 1A). This difference could be due to the different expression of the glycolytic enzymes in different cells (Figure 1B). However, the concentration patterns of the intermediates of different cancer cells are similar between each other. By converting the concentrations of intermediates into Qs and ΔGs of the reactions along the glycolytic flux, we demonstrated that different cancer cell lines shared a same TSAG (Figure 1C & 1D), i.e., Q and ΔG of each reaction in the glycolytic flux were nearly identical in different cells.

**Figure 1.**
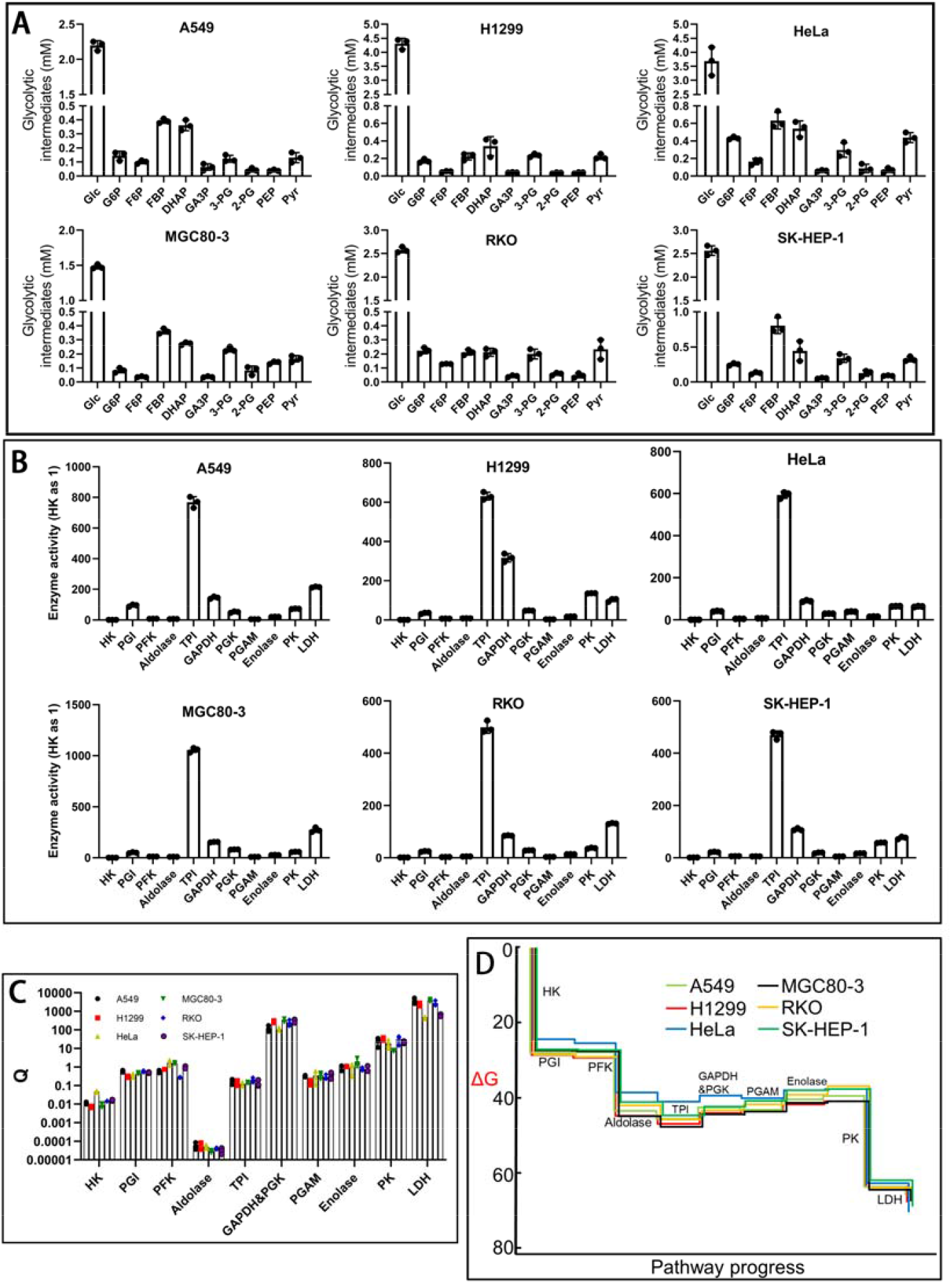
TSAG in cancer cells from different origins. **(A)** Concentrations of glycolytic intermediates (data are from Supplementary Table 1). **(B)** Glycolytic enzyme activity (data from Supplementary Table 2). **(C)** The pattern of glycolytic intermediates in 6 cell lines expressed by Qs (data from Supplementary Table 3). **(D)** ΔG of the reactions along the glycolytic pathway (data from Supplementary Table 3). Data are mean ± SD, n=3. Results were confirmed by two independent experiments.

The reactions catalyzed by PGI, aldolase, GAPDH+PGK, TPI, PGAM, enolase were at near-equilibrium state (Figure 1D), which are important in maintaining the relative concentrations of F6P, G6P, FBP, GA3P, DHAP, 3-PG, 2-PG, and PEP.

The ΔGs for the reactions catalyzed by HK, PFK, and PK were -27.4±1.7, -14.9±1.6, -24.8±1.6, respectively (Figure 1D), generating a major driving force for AG. The ΔG for the reaction catalyzed by LDH was between -2.4±0.4 (MGC80-3) and -7.8±0.4 kJ/mol (HeLa). According to the equation (ΔG = RTln(Q/Keq) = RTln(J-/J+), where J+ and J-refers forward and reverse rate, respectively) (Beard & Qian, 2007; Park et al., 2019), the rate of the forward reactions catalyzed by LDH is 2.6 - 23 folds higher than that of the reverse one, underlying the thermodynamic basis for generating lactate by cancer cells.

Maintaining the near-equilibrium state of the reactions in the flux requires excessive amount of corresponding enzymes. The activities of PGI, aldolase, TPI, GAPDH, PGK1, PGAM, enolase in all the tested cell lines are higher than HK (Figure 1B), the first step of AG, which plays important role in controlling the overall glycolytic rate (Tanner et al., 2018; Wolf et al., 2011). Hence, the activity pattern of glycolytic enzymes underlies the kinetic basis for maintaining near-equilibrium state of these reactions, and it excellently matches TSAG. It is also noted that LDH activity is 63.3±1.9 folds (HeLa) to 272.6±22 folds (MGC80-3) higher than HK (Figure 1B), ensuring a rapid conversion of (pyruvate + NADH) to (lactate + NAD).

Taken together, TSAG is a system property of AG in cancer cells independent of the pathophysiological features of cancer cells. The reactions far from thermodynamic equilibrium state forms the driving force for AG and the reactions at near-equilibrium state plays important role in maintaining the relative concentrations of glycolytic intermediates. These results are consistent with our previous reports (Jin et al., 2020; Xie et al., 2016; Ying, Guo, & Hu, 2019; Zhu et al., 2021).The next question is what is the role of TSAG in AG flux responding to perturbation.

### There is an interrelationship between TSAG, PKM2, and AG flux

PKM2 catalyzes a reaction far from equilibrium state, hence it is classically considered as a preferred site for AG regulation (Anastasiou et al., 2012; Chaneton et al., 2012; Christofk, Vander Heiden, Harris, et al., 2008; Israelsen et al., 2013; Morgan et al., 2013) and perturbing PKM2 is a suitable model to investigate the interrelationship between TSAG, AG flux (rate and intermediate concentration), and PKM2.

At steady-state glycolysis, the rate through PKM2 is determined by the concentration of PKM2, the concentration of its substrates PEP and ADP (because the ΔG of the reactions catalyzed by PKM2 is far from thermodynamic equilibrium, the rate of the reverse reaction could be neglected), and the concentrations of the allosteric effectors in cells. PK rate in the glycolytic flux could be estimated, if these variables could be determined.

To avoid redundancy, PKM2_SA_, PK_SA_, PKM2_AA_, and PK_AA_ refers the specific activity of PKM2 and PK, the actual activity of PKM2 and PK in the glycolytic flux, respectively. For simplicity, the concentration of any compounds would be represented by the symbol [ ], e.g., [G6P] refers the concentration of G6P.

To investigate the interrelationship between PKM2, AG flux, and TSAG in cancer cells, we carried out PKM2 knockdown experiments. PKM2 knockdown significantly reduced PKM2 protein level (Figure 2A). According to Western blotting, the knockdown efficiency varies between 76.7% (HeLa) and 54.2% (RKO). Activity wise, PK_SA_ was reduced by 70% (HeLa) to 30% (RKO). PKM2 knockdown did not significantly affect other glycolytic enzymes (Figure 2B). Thus, PKM2 knockdown only significantly reduced PKM2 without affecting other glycolytic enzymes.

**Figure 2.**
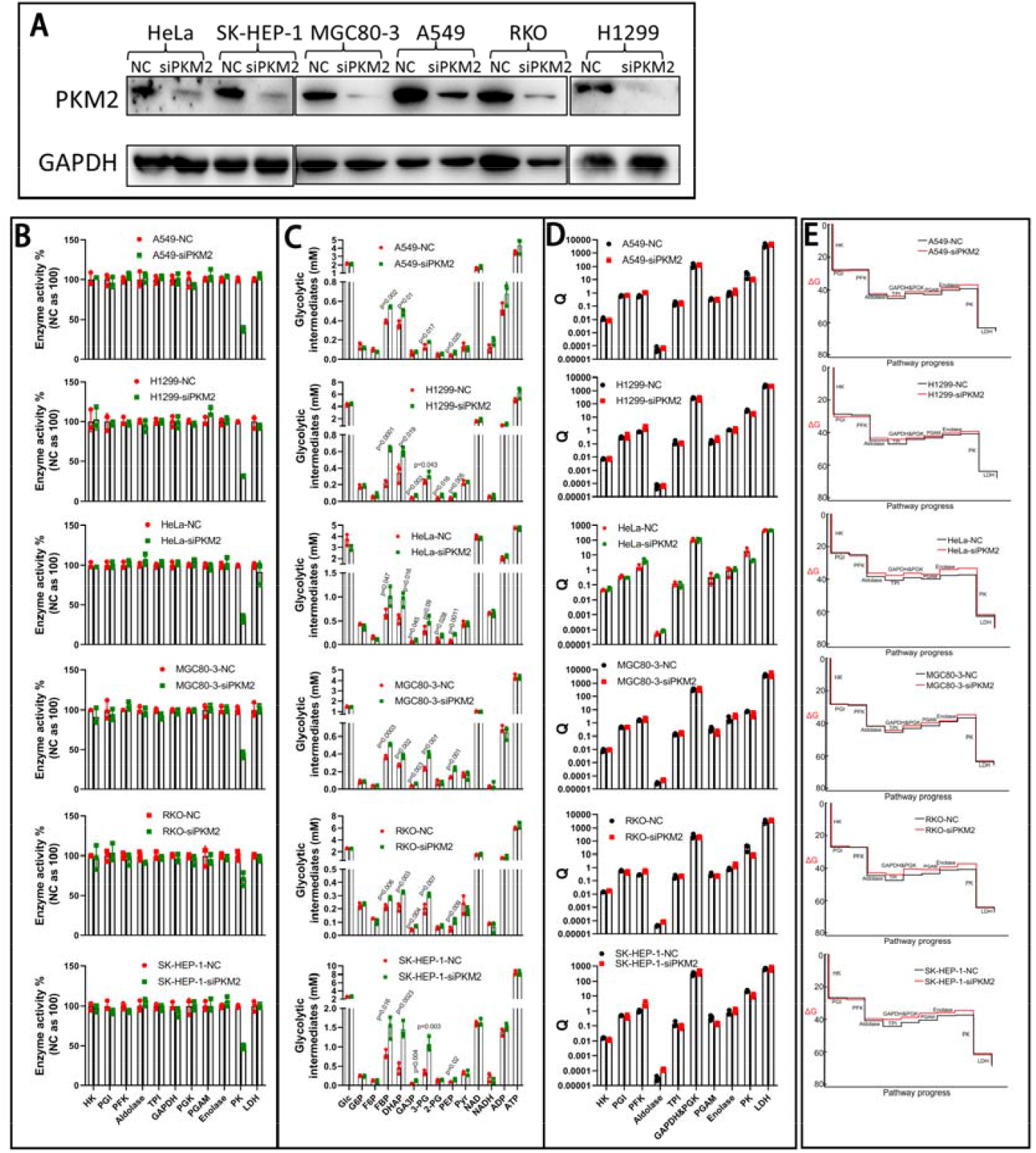
PKM2 knockdown induces a change of the intermediate concentration but does not significantly alter TSAG. **(A)** Western blot of PKM2. **(B)** siRNA knockdown specifically reduced PK activities (data from Supplementary Table 2). **(C)** Glycolytic intermediates (data from Supplementary Table 4). **(D)** Q patterns of the glycolytic intermediates. **(E)** Gibbs free energy of reactions along the glycolytic flux. Data are mean ± SD, n=3. Results were confirmed by two independent experiments.

PKM2 knockdown significantly increased [PEP], [2-PG], [3-PG], [GA3P], [DHAP], and [FBP] lying between PK and PFK1 in the pathway, without significantly affecting [G6P] and [F6P] upstream of PFK1, and [pyruvate] downstream of PKM2 (Figure 2C). [NAD], [NADH], [ADP], and [ATP], which are also intermediates in the glycolytic flux, were comparable between control and PKM2 knockdown cells.

However, the values of Qs (Figure 2D), ΔGs, and ∑ΔGs (Figure 2E) of the glycolytic flux were not significantly affected by PKM2 knockdown. This is consistent in 6 cell lines. The data revealed 2 important points: (1) TSAG in cancer cells is resistant to PKM2 knockdown, and (2) because PKM2-knockdown does not significantly change the ΔG values of the reactions along the glycolytic flux, the concentration change of the glycolytic intermediates responding to PKM2 knockdown, in essence, reflects an equilibration by TSAG. The high activities of aldolase, TPI, GAPDH, PGK1, TPI, PGAM, and enolase rapidly equilibrated [PEP], [2-PG], [3-PG], [GA3P], [DHAP], and [FBP] in both control and PKM2-knockdown cells.

The ΔG of PFK1-catalyzed reaction was -14.0±1.2 kJ/mol in PKM2-knockdown cells (Figure 2E), not significanlty different from that in conotrol cells. This thermodynamic barrier explains why PKM2 knockdown cannot affect [G6P] and [F6P]. Pyruvate concentrations between PKM2-knockdown and control cells were comparable without significant difference, suggesting that PK_AA_ in AG flux was not significantly affected by PKM2 knockdown.

Finally, AG rate (glucose consumption rate and lactate generration rate) was not significantly affected by PKM2 knockdown in all the tested cell lines (Figure 3), suggesting that PK_AA_ in control cells and PKM2-knockdown cells were not significantly different from each other. To further validate lactate generated by the cancer cells is indeed from glucose, we used [^13^C_6_]-glucose tracing, which confirmed that lactate-m3% were comparable between control and PKM2-knockdown cells (Supplementary Figure 1). The results thus point out that the rates of glucose carbon through PKM2 between control and PKM2-knockdown cells are not significantly different from each other.

**Figure 3.**
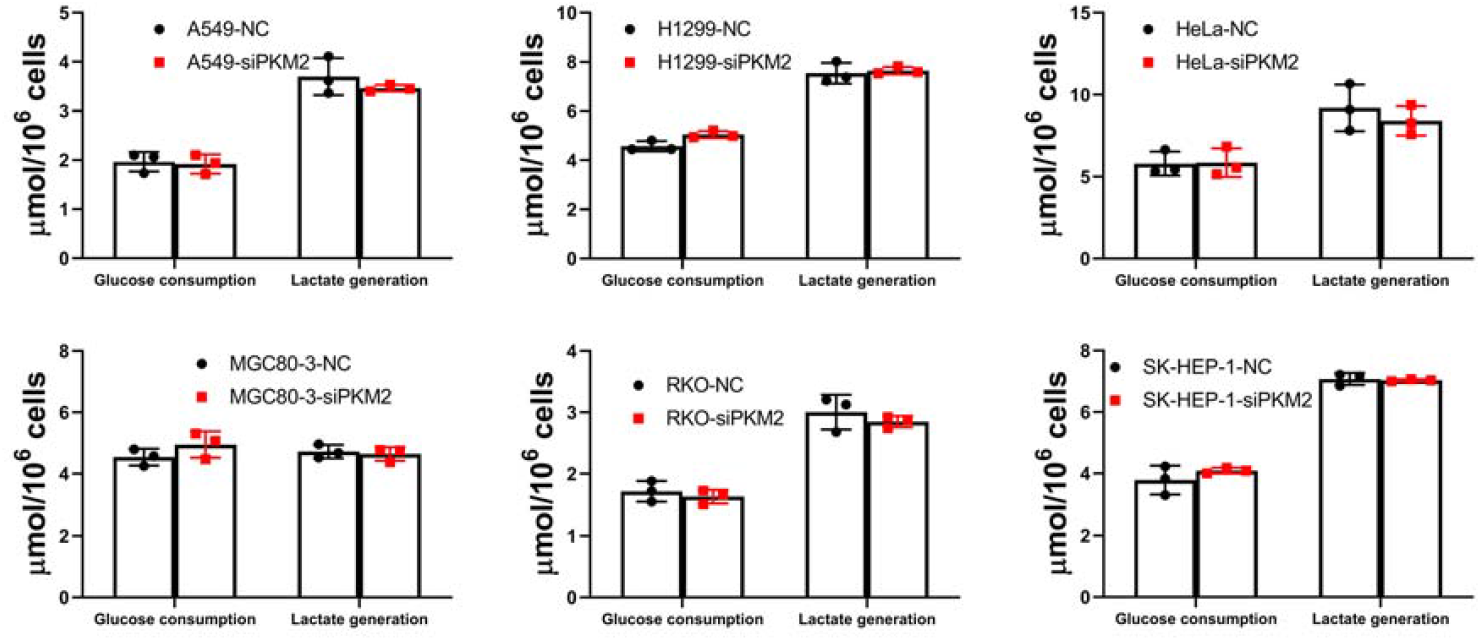
PKM2 knockdown does not significantly inhibit glucose consumption and lactate generatoin by cancer cells. Cells were transfected with NC or siPKM2 for 48 hours, cultured in fresh complete medium with 10% FBS for 6 hour, and culture medium were collected for glucose consumption and lactate generation determination. Data are mean ± SD, n=3. Results were repeated by two independent experiments.

Collectively, PKM2 knockdown significantly reduced PK_SA_ but did not significantly affect AG rate, revealing that PK_AA_ is not significantly perturbed; PKM2 knockdown induced a significant increase of [PEP], [2-PG], [3-PG], [GA3P], [DHAP], and [FBP], but did not alter the ΔGs of the reactions along pathway, indicating that the concentration increase of the intermediates responding to PKM2 knockdown reflects an equilibration by TSAG. The next question is to investigate the interrelationship between AG rate, PK_AA_, and TSAG in the flux, which may interpret why PKM2 knockdown does not influence PK_AA_ in the flux.

### TSAG controls reciprocal [PEP] and [PKM2]

The data in Figure 2 tell that there is a reciprocal relationship between [PKM2] and [PEP] in AG flux: in control cells, [PKM2] is high and [PEP] is low, while in PKM2 knockdown cells, [PKM2] is low and [PEP] is high. Because PK_AA_ in the glycolytic flux is a function of both [PKM2] (or PK_SA_) and [PEP] (here [ADP] is not taken into consideration, because it remains unchanged). PK-catalyzed reaction in AG flux is far from equilibrium, e.g., the ΔGs of this reaction in control and PKM2-knockdown cells (HeLa) were -25.4 and -28.6 kJ/mol, so that the reverse reaction rate could be neglected. The reasoning suggests that the reciprocal relationship between [PKM2] and [PEP] may keep a constant PK_AA_ in the glycolytic flux. If so, TSAG, [PKM2], [PEP], PK_AA_, and AG flux could be rationally interconnected. However, the allostery of PKM2 must also be taken into consideration.

### FBP dominates allostery of PKM2

PKM2 was regulated by multiple allosteric effectors in cells, including but not limited to inhibitors (alanine, phenylalanine, proline, tryptophan, valine) and activators (FBP, serine) (Chaneton et al., 2012; Macpherson et al., 2019; Morgan et al., 2013; Yuan et al., 2018). PEP concentrations in these cells are between 0.1-0.2 mM(Jin et al., 2020; Zhu et al., 2021) (Figure 2). As it is more meaningful to measure the allosteric regulation of PKM2 at cellular or physiologically relevant concentrations, we used 0.2 mM PEP in the assay for measuring PK allostery.

The sensitivity of PK from different cancer cells to allosteric inhibitors differed from each other, as manifested by the IC_50_s (Supplementary Figure 2A, Supplementary Table 5). AC_50_s of PK from different cells to FBP varies from 22.4±14.3 (SK-HEP-1) to 57.5±22.7 (RKO) nanomolar range (Supplementary Figure 2B, Supplementary Table 5), whereas AC_50_s to serine and IC_50_s to inhibitors were in micro molar range (Supplementary Figure 2C, Supplementary Table 5). When FBP saturates PK, the maximal activity of PK from 5 cancer cell lines increased by 3.3 folds (HeLa) and 1.9 folds (RKO) (Supplementary Figure 2B).

For PK in cell lysate, alanine was the most potent inhibitor, followed by phenylalanine, tryptophan, valine, and proline, consistent in 5 cancer cell lines (Supplementary Figure 2A, Supplementary Table 5). For pure recombinant PKM2, phenylalanine was the most potent inhibitor (Supplementary Figure 2A, Supplementary Table 5), consistent with the previous report (Morgan et al., 2013).

PK in cell lysate was in general more sensitive to allosteric inhibitors than pure recombinant PKM2 (Supplementary Figure 2A). 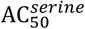 of PK in cell lysate is markedly lower than that of pure recombinant PKM2 (Supplementary Figure 2C, Supplementary Table 5). Although 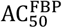 of PK in cell lysate and recombinant PKM2 were comparable (Supplementary Figure 2B, Supplementary Table 5), the maximal activity of the latter activated by FBP was markedly higher than the former (Supplementary Figure 2B).

Together, the results showed that PKs in different cells are all allosteric enzymes, although they also exhibit some differences.

In order to study the combined effect of these regulators on PK, we determined these regulators in 5 cancer cell lines (Table 1). Among the 5 allosteric inhibitors, the concentration of alanine was the highest in average and the concentration of tryptophan was the lowest in average. The concentrations of these regulators in different cells varied from cell to cell. The concentration of alanine ranged between 1.36±0.50 mM (HeLa) and 5.78±1.57 mM (RKO). According to the curves of PK activity versus alanine (Supplementary Figure 2), we could estimate PK activity exposed to such alanine concentration in cells. In the 5 cancer cells, alanine alone could inhibit PK by 89% (MGC80-3) to 98% (SK-HEP-1) (Table 1). In the same way, we could estimate PK activity according to the cellular concentration of each allosteric inhibitor or activator (Table 1).

**Table 1.** Cellular concentrations (mM) of allosteric regulators and the corresponding PK activity (%). Data represent the mean ± SD of 3 independent experiments with each experiment performed in triplicates.

|  | Ala | PK activity* | Phe | PK activity* | Pro | PK activity* | Trp | PK activity* | Val | PK activity* | FBP | PK activity* | Ser | PK activity* |
| --- | --- | --- | --- | --- | --- | --- | --- | --- | --- | --- | --- | --- | --- | --- |
| A549 | 1.53±0.30 | 2.03±0.21 | 0.20±0.06 | 11.22±1.32 | 0.50±0.07 | 38.34±3.40 | 0.14±0.13 | 49.63±23.98 | 0.51±0.11 | 28.70±4.73 | 0.14±0.03 | 266.5±0.44 | 1.61±0.35 | 250.75±1.05 |
| HeLa | 1.36±0.50 | 5.35±0.11 | 0.29±0.04 | 8.60±0.48 | 1.39±0.15 | 9.22±1.25 | 0.12±0.01 | 40.79±2.71 | 0.70±0.21 | 15.81±4.56 | 0.38±0.04 | 325.5±0.15 | 1.39±0.34 | 254.86±1.74 |
| MGC80-3 | 2.15±0.87 | 11.16±0.42 | 0.26±0.03 | 14.04±0.36 | 1.05±0.10 | 23.49±1.21 | 0.19±0.15 | 47.53±15.53 | 0.78±0.19 | 25.46±3.62 | 0.28±0.01 | 271.2±0.04 | 1.46±0.08 | 248.34±0.06 |
| RKO | 5.78±1.57 | 5.65±0.03 | 0.33±0.05 | 11.49±0.57 | 3.16±0.52 | 6.65±1.66 | 0.22±0.24 | 52.25±26.17 | 0.92±0.32 | 20.66±9.11 | 0.37±0.03 | 217.3±0.00 | 2.79±0.41 | 186.11±0.10 |
| SK-HEP-1 | 3.02±0.15 | 1.51±0.02 | 0.40±0.06 | 4.58±0.73 | 1.98±0.47 | 9.55±3.48 | 0.19±0.15 | 43.00±19.34 | 0.96±0.32 | 10.80±4.39 | 0.24±0.04 | 286.2±0.00 | 2.60±0.71 | 257.56±0.37 |

\*The corresponding PK activity (% of no inhibition) to each of the allosteric regulator is estimated according to the kinetic curves in Supplementary Figure 2.

In cancer cells, PKM2 is co-regulated by numerous allosteric effectors, the concentrations of these allosteric effectors could change dynamically, hence it is generally conceived that PKM2 is dynamically equilibrated between different states, for example, T and R state, tetramer and dimer, in cancer cells, enabling highly dynamic change of PKM2 activity(Chaneton et al., 2012; Christofk, Vander Heiden, Harris, et al., 2008; Christofk, Vander Heiden, Wu, Asara, & Cantley, 2008; Israelsen et al., 2013; Keller, Doctor, Dwyer, & Lee, 2014; Morgan et al., 2013; Yuan et al., 2018). Nevertheless, PK activity exposed to the mixture of allosteric activators and inhibitors at their cellular concentrations is not determined in the previous studies.

We then assayed PK activity in the cell lysate in the presence of these allosteric regulators at their cellular concentrations, for example, in HeLa cells, the concentrations of alanine, phenylalanine, proline, tryptophan, valine, serine, and FBP were 1.36±0.50, 0.29±0.04, 1.39±0.15, 0.12±0.01, 0.70±0.21, 1.39±0.34, and 0.38±0.04 mM, respectively, so when determining PK activity in the HeLa cell lysate, the assay mixture contained these regulators at the above indicated concentrations, and so forth for other cells. Unless otherwise stated, the cellular concentrations of FBP, serine, and amino acid mixture in the subsequent Figures and text are the same as above.

PK activity was enhanced by serine and FBP in the absence of allosteric inhibitor mixture (alanine, phenylalanine, proline, tryptophan, valine) (Figure 4), which is termed AIM in the manuscript. The potency of serine and FBP to activate PK activity was comparable with each other.

**Figure 4.**
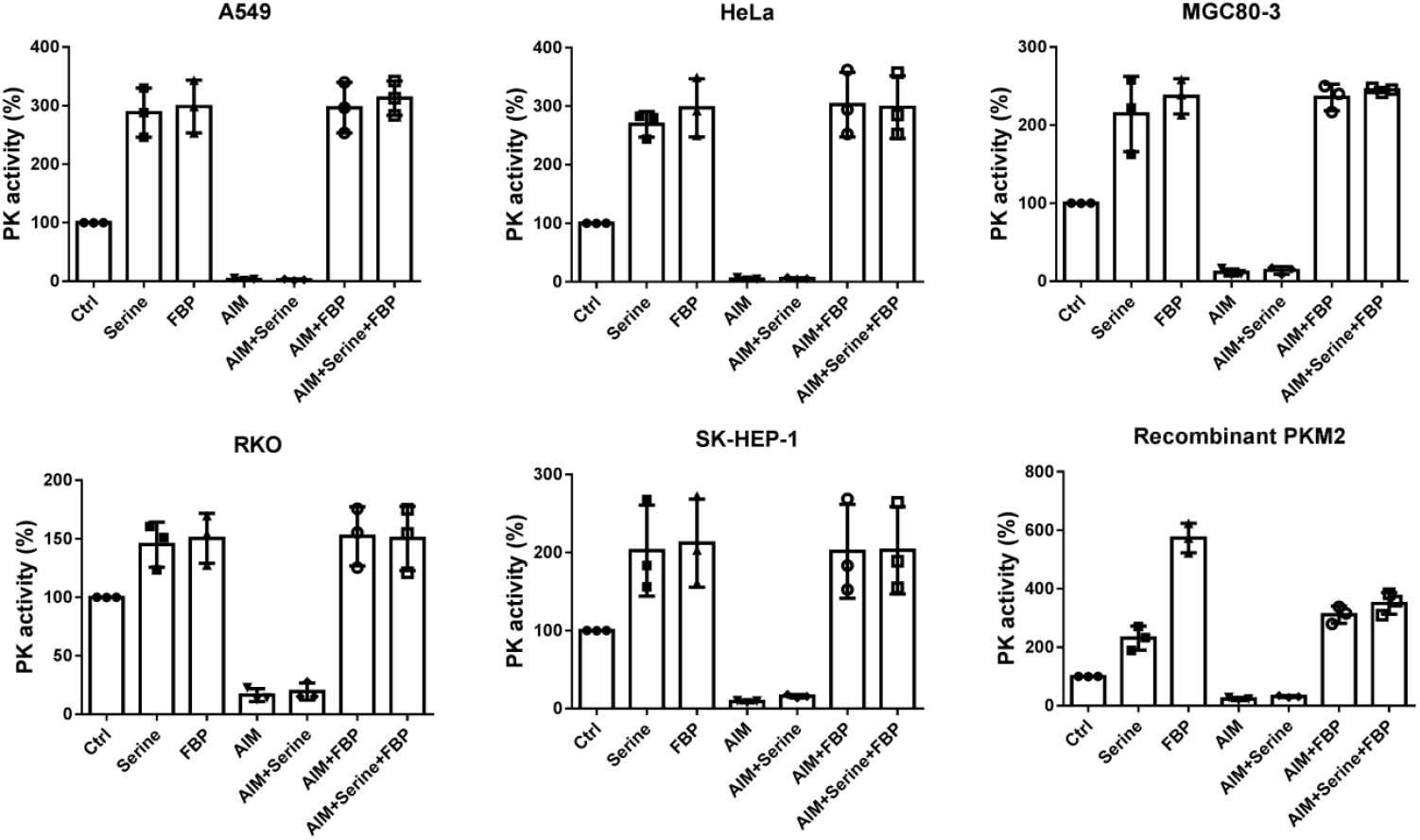
The dominating role of FBP in the allosteric regulation of cell lysate PK or recombinant PKM2. For cell lysate, cellular concentration of FBP, serine, and AIM were used. For pure recombinant PKM2, AIM used were 2.0 mM alanine, 2.0 mM phenylalanine, 5.0 mM proline, 2.5 mM tryptophan and 3.0 mM valine; Serine and FBP concentrations used were 5 mM and 0.1 mM, respectively. The substrate concentrations were 2 mM ADP and 0.2 mM PEP. PK activities assayed with no inhibitors or activators were set as control (100%). Data represent the mean ± SD of 3 independent experiments with each experiment performed in triplicates.

When PK activity was assayed with AIM without serine or FBP, the residual PK activities were in the range between (16±5.5) % (RKO) and (3.0±1.9) % (A549) (Figure 4). Addition of serine did not significantly reverse the inhibition. Addition of FBP not only completely reversed the inhibition but also activated PK activity to the level without AIM. The effect of FBP plus serine on PK inhibition by AIM was the same as that of FBP alone (Figure 4).

For pure recombinant PKM2 (Figure 4), in the absence of AIM, serine and FBP could activate PKM2 activity by 2.3±0.4 and 5.7±0.5 folds, respectively, indicating FBP is significantly more potent than serine. In the presence of AIM, the residual PKM2 activity was (23±5.3) %. Serine did not significantly reverse AIM-induced inhibition of PK, whereas FBP could reverse AIM-induced inhibition of PKM2, but could not enhance its activity to the level without AIM.

Further, we sought to determine the 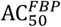 and 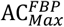 for PK with or without saturating concentration of AIM. The concentration of each allosteric inhibitor in this assay is about 10 folds of the IC_50_ (2.0 mM alanine, 0.7 mM phenylalanine, 2 mM proline, 1 mM tryptophan and 1.0 mM valine for cell lysate; 2.0 mM alanine, 0.7 mM phenylalanine, 5 mM proline, 2.5 mM tryptophan and 3.0 mM valine for pure recombinant PKM2). According to the curves (Figure inhibition of PK by the AIM (Figure 5, Table 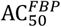 is 28.9±21.0 nM (SK-HEP-1) to 67.2±32.4 nM (RKO), the minimum concentration for maximal activation is 216.4±52.8 nM (HeLa) to 279.0±143.1 nM (A549) (Table 2). With AIM, 349.6±31.2 nM (MGC80-3) to 585.2±99.0 nM (HeLa) FBP could completely reverse the inhibition of PK by the AIM (Figure 5, Table 2), 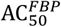 were 682.1±126.2 nM (A549) to 1066.1±393.6 nM (RKO), 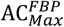 was 1258.9±245.4 nM (A459) to 2293.7±503.7 nM (HeLa). Note that the maximal activity of PK activated by FBP with or without AIM were comparable without statistical significance (Figure 5).

**Figure 5.**
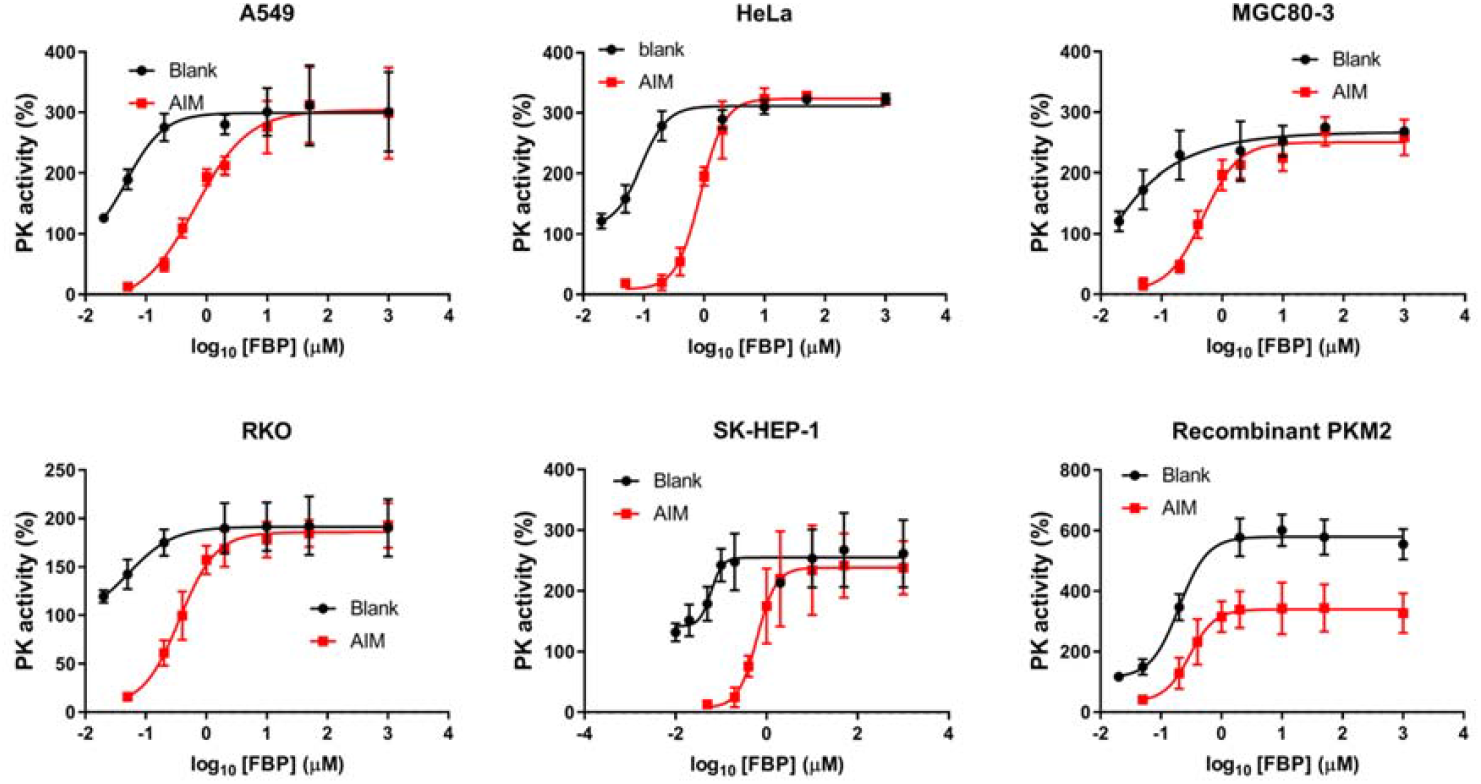
The dominating role of FBP in the allosteric activation of PK. Kinetic profiles of pyruvate kinase activities at different FBP concentrations with or without AIM. For cell lysate, AIM was consisted of 2.0 mM alanine, 0.7 mM phenylalanine, 2 mM proline, 1 mM tryptophan and 1.0 mM valine. For pure recombinant PKM2, AIM used was 2.0 mM alanine, 2.0 mM phenylalanine, 5.0 mM proline, 2.5 mM tryptophan and 3.0 mM valine. The substrate concentrations were 2 mM ADP and 0.2 mM PEP. PK activities measured without FBP or AIM were used as control (100%). Data represent the mean ± SD of 3 independent experiments with each experiment performed in triplicates.

**Table 2.** 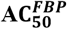 and 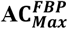 (nM) for cell lyate PK or recombinant PKM2 in the absence or in the presence of cellular AIM. Data are estimated from the kinetic curves in Figure 5 and represent the mean ± SD of 3 independent experiments with each experiment performed in triplicates.

|  | A549 |  | HeLa |  | MGC80-3 |  | RKO |  | SK-HEP-1 |  | Recombinant PKM2 |  |
| --- | --- | --- | --- | --- | --- | --- | --- | --- | --- | --- | --- | --- |
|  | Control | AIM | Control | AIM | Control | AIM | Control | AIM | Control | AIM | Control | AIM |
| $AC_0^*$ | / | 377.9±35.1 | / | 585.2±99.0 | / | 349.6±31.2 | / | 394.2±121.8 | / | 571.2±132.1 | / | 170.1±70.2 |
| $AC_{50}^{FBP}$ | 33.2±11.2 | 682.1±126.2 | 43.5±18.6 | 813.7±41.8 | 43.9±18.4 | 698.5±313.1 | 67.2±32.4 | 1066.1±393.6 | 28.9±21.0 | 870.2±345.4 | 52.3±17.3 | 246.4±85.0 |
| $AC_{Max}^{FBP}$ | 279.0±143.1 | 1258.9±245.4 | 216.4±52.8 | 2293.7±503.7 | 252.2±133.6 | 1624.1±611.9 | 273.3±129.3 | 1518.4±359.0 | 231.5±128.3 | 1747.5±588.0 | 731.5±169.8 | 1190.6±208.3 |

\* denotes the concentration of FBP (nM) that completely reverse the inhibition of PK by AIM, i.e., the activity is equal to control without AIM.

For pure recombinant PKM2, without AIM, the 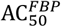 is 52.3±17.3 nM, 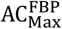 is 731.5±169.8 nM. In the presence of AIM, 170 nM FBP completely reversed the inhibition of PK by the AIM, 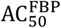 and 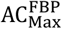 were 246.4±85.0 and 1190.6±208.3 nM, respectively (Figure 5, Table 2). Note that the maximal activity of PKM2 in the absence of AIM was 1.8 folds higher than that in the presence of AIM (Figure 5).

Collectively, in the co-presence of AIM and serine, AIM dominates the allosteric inhibition of cell lysate PK and recombinant PKM2; in the co-presence of AIM and FBP, FBP dominates the allosteric activation of cell lysate PK and recombinant PKM2. This is consistent with the previous reports that inhibitory amino acids fail to inhibit PKM2 in the presence of FBP, indicating the allosteric dominance of FBP (Macpherson et al., 2019; Morgan et al., 2013; Sparmann, Schulz, & Hofmann, 1973; Yuan et al., 2018).

### TSAG stabilizing PK_AA_ by tuning the reciprocal [PKM2] and [PEP]

Cellular [FBP] in control and PKM2 knockdown cells (Figure 2) are far higher than 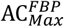 (Table 2), supporting that the allosteric regulation of PK in cancer cells is dominated by FBP. As such, the allostery of PK in cancer cells could be considered as a constant when taking it into calculating PK_AA_ in cancer cells.

To this stage, we could measure PK_AA_ in control and PKM2 knockdown cells using biochemical approaches. PK_AA_ in cell lysate prepared from PKM2-knockdown cells and control cells was assayed at cellular concentrations of PEP, with or without allosteric effectors also at their cellular concentrations (Figure 6). Without allosteric regulators, PK_AA_ were comparable with or without PKM2 knockdown in A549, H1299, and MGC80-3 cells, but it was significantly higher in PKM2-knockdown cells than control cells (RKO, SK-HEP-1, and HeLa). With allosteric regulators (FBP, serine, and AIM at their cellular concentrations), PK_AA_ was comparable with or without PKM2-knockdown in all the tested cells (Figure 6).

**Figure 6.**
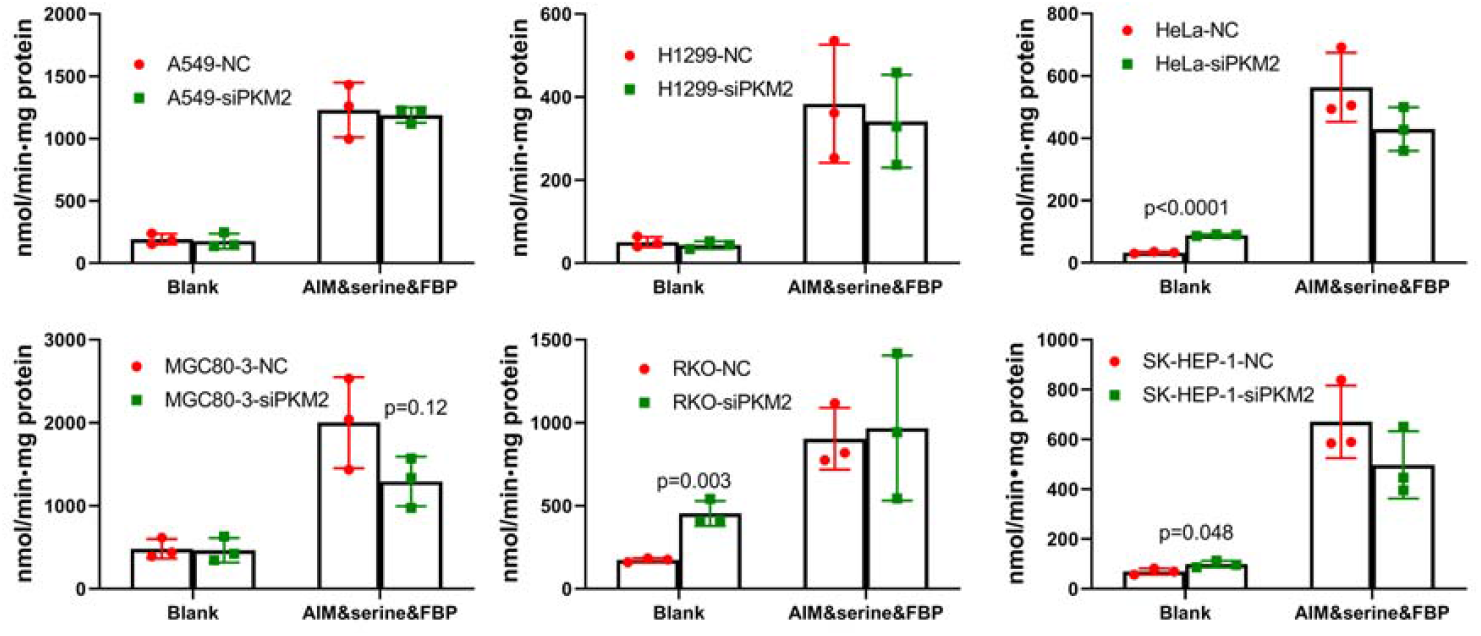
PK_AA_ in control and PKM2 knockdown cells. PK activities were measured at cellular concentration of PEP in the presence or absence of allosteric regulators (AIM, seine, FBP) at their cellular concentrations. Data represent the mean ± SD of 3 independent experiments with each experiment performed in triplicates.

Because TSAG is essential for maintaining [FBP] as well as the reciprocal relationship between [PEP] and [PKM2] in AG flux, it plays an important role in stabilizing PK rate in AG flux.

### The results of the PKM2 knockout experiment are consistent with those of the PKM2 knockdown

PKM2 knockout in Hela cells resulted in 83% decrease in PK_SA_ by activity assay and invisible PKM2 bands by Western blot (Figure 7A). However, it did not significantly affect glucose consumption. Lactate production was significantly reduced by PKM2 knockout, but only by 13% in absolute value (Figure 7B).

**Figure 7.**
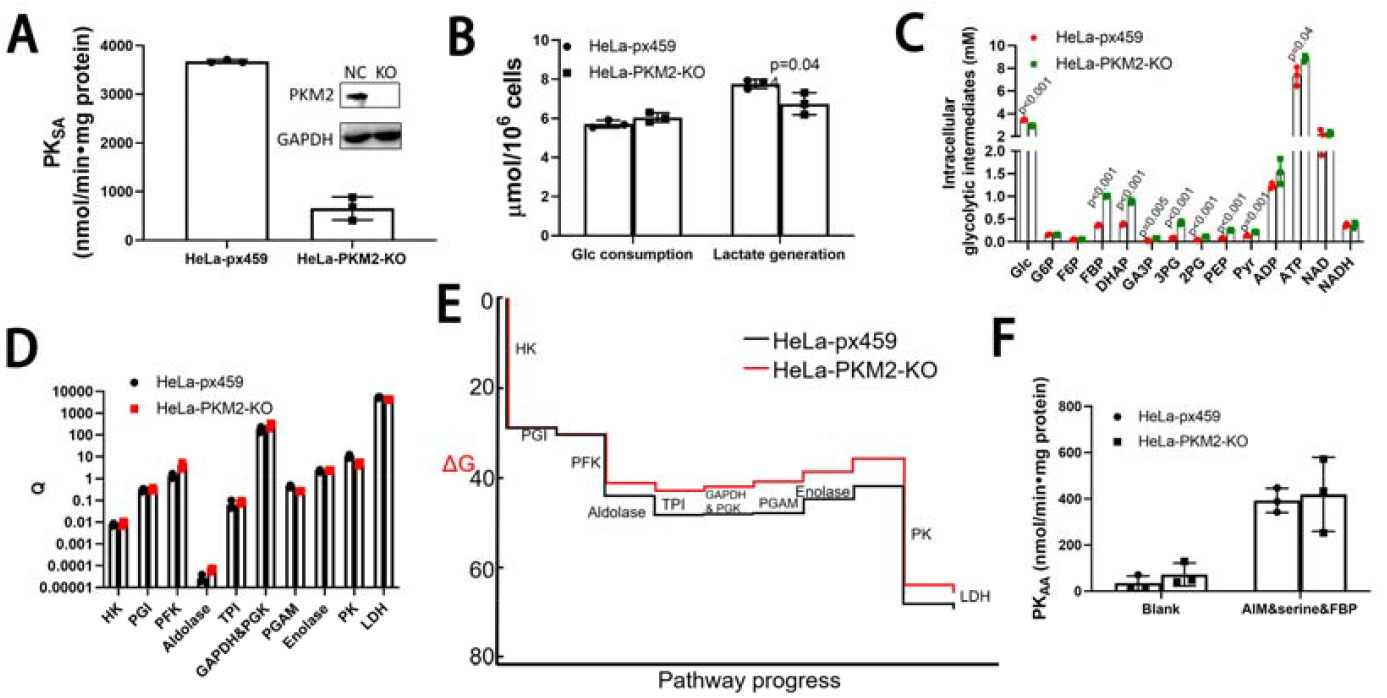
AG rate, TSAG, and PK_AA_ in HeLa cells with or without PKM2 knockout. **(A)** Validation of PKM2 knockout by enzyme activity assay and western blot. **(B)** Glucose consumption and lactate generation in control and PKM2 knockout cells. **(C)** Glycolytic intermediates. **(D)** Q pattern of glycolytic intermediates. **(E)** Gibbs free energy of reactions along the glycolytic flux. **(F)** PK_AA_ in control and PKM2 knockout cell lysate. PK activities were measured at cellular concentration of PEP in the presence or absence of allosteric regulators (AIM, seine, FBP) at their cellular concentrations. Data are mean ± SD, n=3. Results were repeated by 3 independent experiments.

Similar to PKM2 knockdown, PKM2 knockout significantly increased [FBP], [DHAP], [GA3P], [3-PG], [2-PG], [PEP] in AG flux (Figure 7C), but did not significantly change the Q and ΔG of the corresponding reactions in the flux (Figure 7D&E). These results demonstrated that PKM2 knockout-induced concentration changes of the intermediates in AG flux were controlled by TSAG.

Finally, we assayed PK_AA_ in control and PKM2 knockout cell lysate. The enzymatic activity was measured according to cellular [PEP], [FBP], and [AIM]. With or without allosteric modulators, there was no significant difference of PK_AA_ between control and PKM2 knockout cell lysate(Figure 7F). The result is consistent with that of the PKM2 knockdown that TSAG stabilizes PK_AA_ in AG flux by regulating the relative concentrations of [PEP] and [PK]. The residual activity of PK left in PKM2 knockout cells should be derived from PKL/R. These PK isoenzymes are also expressed in cancer cells and are also allosteric enzymes (Llorente, Marco, & Sols, 1970; Yamada & Noguchi, 1999).

### PKM2 knockdown moderately enhances glucose carbon to serine synthesis pathway

In PKM2 knockdown cells, a higher and stable [3-PG] is maintained by TSAG. Because 3-PG is the precursor of serine, glucose carbon to serine synthesis may increase. We traced [^13^C_6_]Glc to serine (Figure 8). The m0 species of serine and m0 glycine are provided by the culture medium, and the m3 serine and m2 glycine are generated from [^13^C_6_]Glc through 3-PG. Knockdown of PKM2 did not affect the glycolysis rate within 9 hours (Figure 8A), consistent with the results in Figure 3. HeLa-siPKM2 showed a higher rate of both m0 serine consumption and m3 serine generation than HeLa-NC (Figure 8B). Serine could be converted to glycine via serine hydroxymethyl transferase in cells. In consistency with serine metabolism, HeLa-siPKM2 showed a higher rate of both m0 glycine consumption and m2 glycine generation than HeLa-NC (Figure 8C).

**Figure 8.**
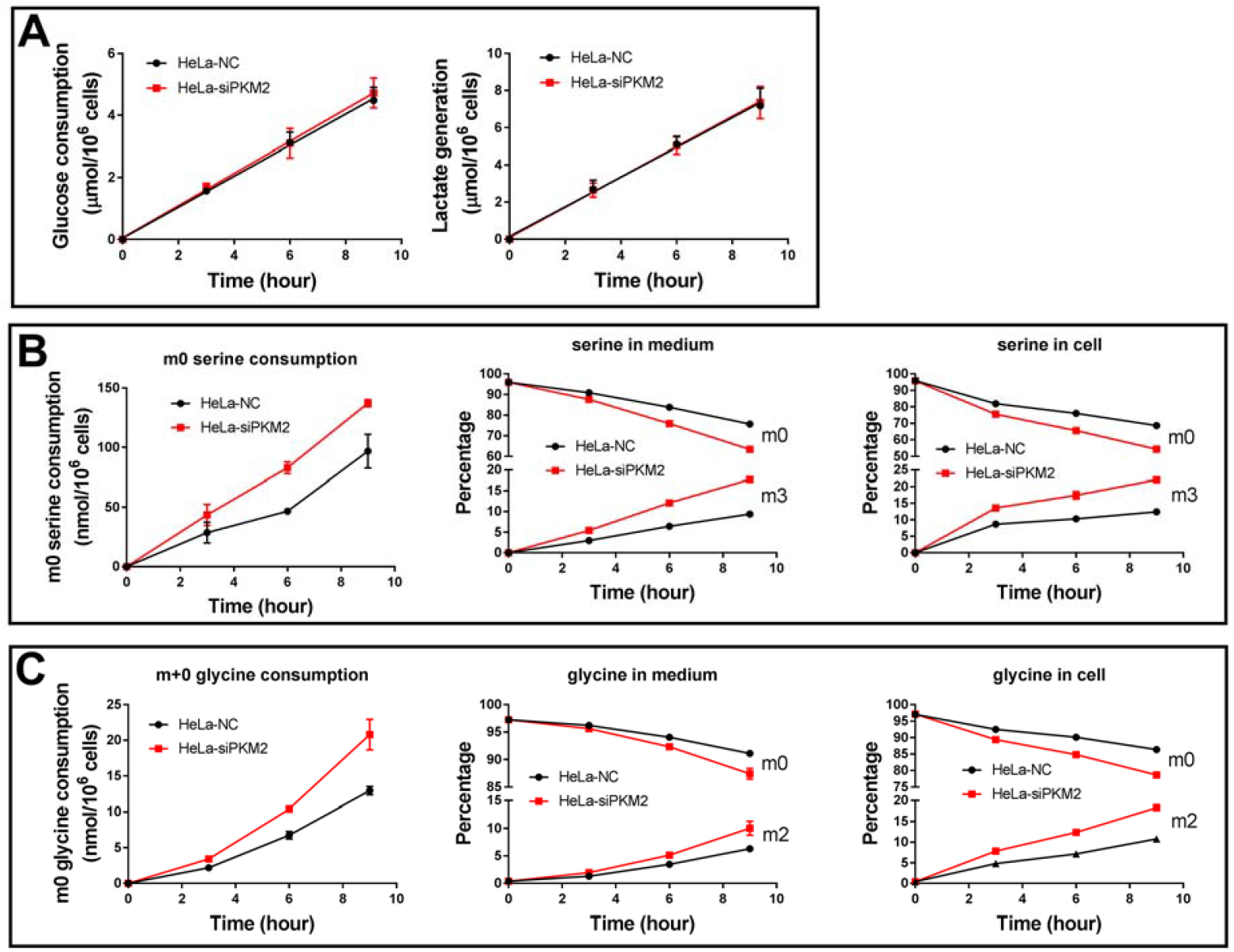
PKM2 knockdown moderately but significantly increases glucose carbon to serine synthesis pathway. **(A)** Glucose consumption and lactate generation by HeLa-NC and HeLa-siPKM2 cells. **(B)** Serine consumption and serine synthesis by HeLa-NC and HeLa-siPKM2. Left panel, consumption of m0 serine provided by the culture medium; middle panel, percentage of m0 and m3 serine in medium; right panel, percentage of m0 and m3 serine in cells. **(C)** Glycine consumption and glycine synthesis by HeLa-NC and HeLa-siPKM2 cells. Left panel, consumption of m0 glycine provided by the culture medium; middle panel, percentage of m0 and m2 glycine in medium; right panel, percentage of m0 and m2 glycine in cells. Data represent the mean ± SD of 3 independent experiments with each experiment performed in triplicates.

### PKM2 knockdown only exerts a marginal effect on cell growth rate

HeLa-siPKM2 harbored a marginally but significantly higher percentage of S phase cells, lower percentage of G0&G1 phase cells, and comparable percentage of G2/M phase cells, in comparison to HeLa-NC (Figure 9A & B). HeLa-siPKM2 also showed a marginally but not significantly higher growth than HeLa-NC (Figure 9C), accompanied with a marginally higher glucose consumption and lactate generation (Figure 9D).

**Figure 9.**
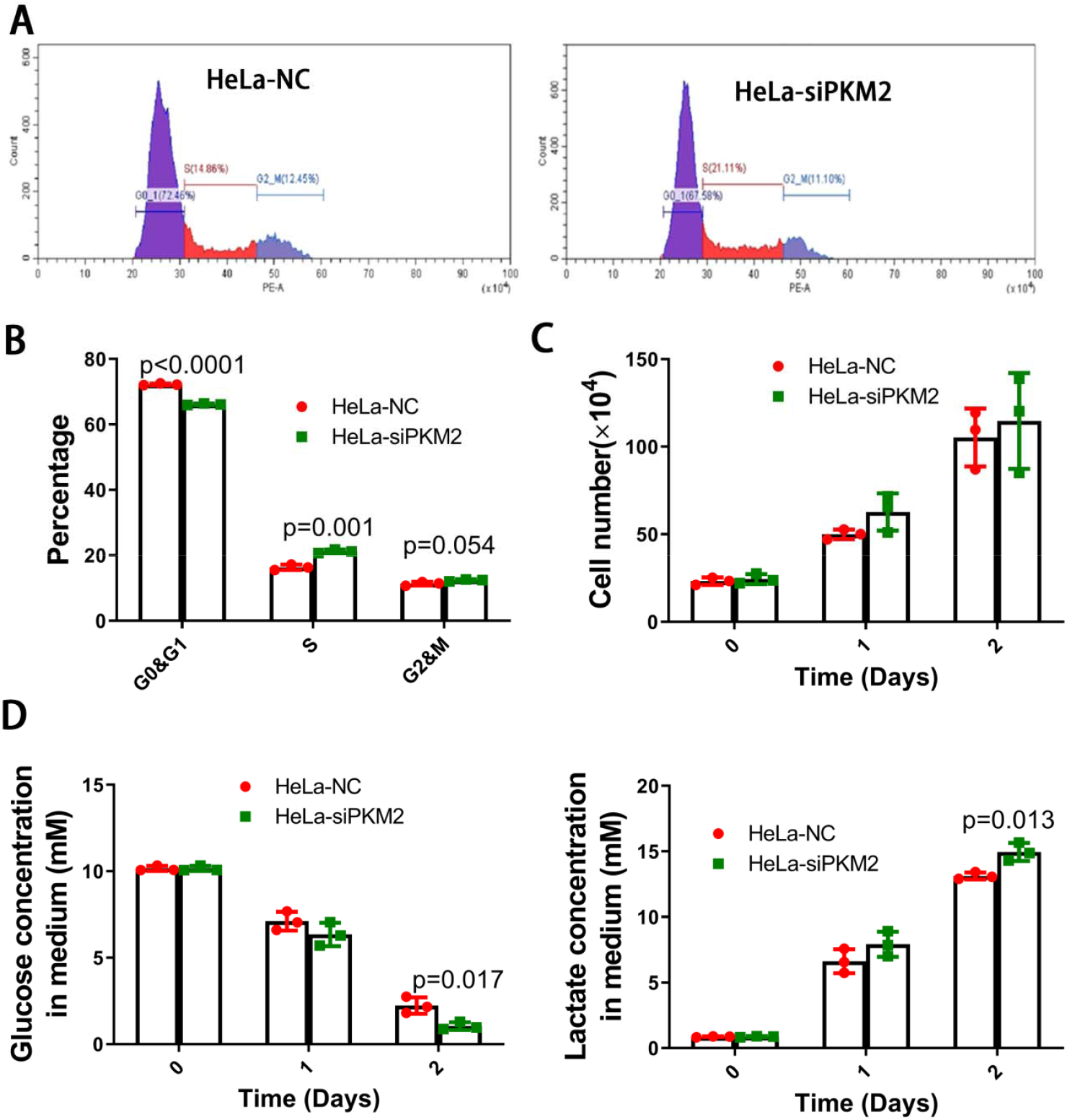
Effect of PKM2 knockdown on cell growth. **(A)** Representative flow cytometry images of cell cycle analysis of HeLa-NC and HeLa-siPKM2 cells. **(B) C**ell cycle distribution of HeLa-NC and HeLa-siPKM2 cells. **(C) C**ell number of HeLa-NC and HeLa-siPKM2 cells at 0, 1, and 2 days after the transfected cells were seeded to new plates. **(D)** Glucose and lactate concentrations in culture medium of HeLa-NC and HeLa-siPKM2 cells. Data represent the mean ± SD of 3 independent experiments with each experiment performed in triplicates.

### Thermodynamic state of cell-free glycolysis system is similar to TSAG in cells

In cells, glycolysis is connected to its subsidiary pathways and it is integrated into a whole network of metabolism. It is not known if the metabolic network influences the thermodynamic state of the glycolytic flux. To investigate this issue, we used a cell-free glycolysis, which could be considered as an isolated glycolysis system (Jin et al., 2020; Xie et al., 2016; Zhu et al., 2021).

The intermediate concentration in cell-free system is 1 order of magnitude lower than that in living cells (Figure 10A, Supplementary Table 1&6). This is most probably because enzyme concentrations in living cells are far higher than that in cell-free system. However, ΔGs of the reactions along the glycolytic flux were similar between cell free and living cell glycolysis (Figure 10B & 2E).

**Figure 10.**
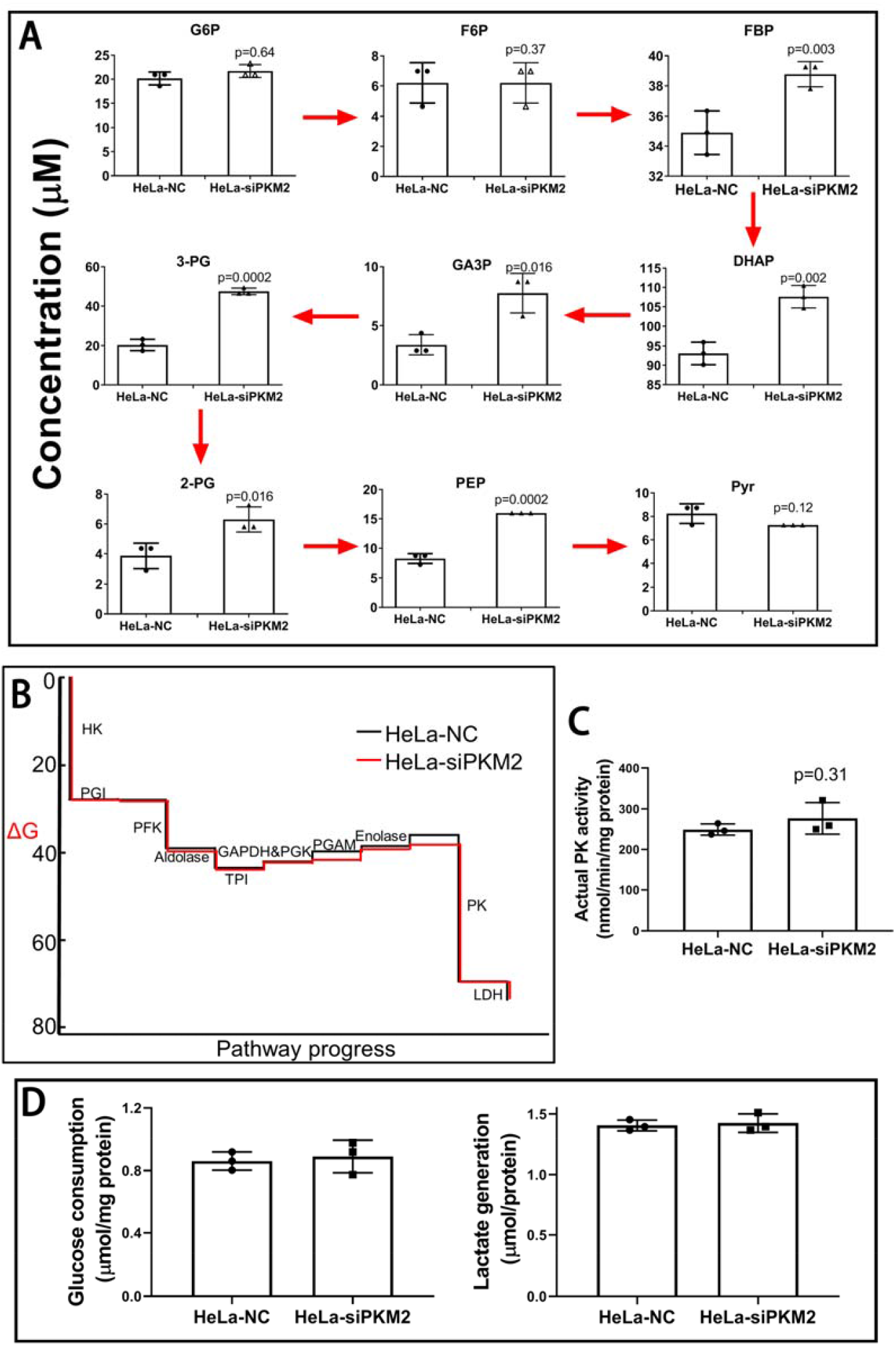
Cell-free glycolysis model. **(A) C**oncentrations of glycolytic intermediates in the reaction mixture. PK of HeLa cells were knocked down by siPKM2, and both control and PKM2 knockdown cells were collected and lysed. The cell lysate was added into reaction buffer to start the glycolysis reaction. Glucose, lactate and glycolytic intermediates were measured. **(B) C**umulative ΔG of the reactions along the glycolysis. **(C)** Actual PK activity determined according to PEP and FBP concentrations in A. **(D)** Glucose consumption rate and lactate generation rate in the reaction mixture. Data are mean ± SD, n=3. Results were repeated by two independent experiments.

To investigate if thermodynamic state of the cell-free glycolytic flux is resistant to PKM2 perturbation, we used cell lysate prepared from control and PKM2 knockdown cells. PK_SA_ in PKM2 knockdown cells is 30% of that in control cells. [G6P], [F6P], [pyruvate] were not significantly different from each other, the concentrations of glycolytic intermediates between PFK and PK were significantly increased (Figure 10A) in HeLa-siPKM2 lysate group, and the ΔGs of these reactions were comparable with each other (Figure 10B).

To investigate if thermodynamic state also stabilizes PK_AA_ in the cell-free glycolysis, we used cell lysate prepared from control and PKM2 knockdown cells. [FBP] generated from glycolytic flux in PKM2 knockdown and control group both saturated PKM2 (Figure 10B). [PEP] in PKM2 knockdown and control group was 16.0 μM and 8.2 μM, respectively. By taking [FBP], [PEP], and [PKM2] into consideration, there was no significant difference of PK_AA_ between 2 groups (Figure 10C), despite a marked difference of PK_SA_. Consistently, glucose consumption and lactate generation were comparable between the 2 groups (Figure 10D).

Taken together, the thermodynamic state of cell-free glycolytic system shares the same feature of TSAG in living cells, and it plays an important role in stabilizing PK_AA_ by tuning reciprocal relationship between [PKM2] and [PEP] in the flux.

## Discussion

Our work demonstrated that TSAG is nearly identical in cancer cells from different origins and cancer types hence it is a system property of AG independent of the pathophysiological features of cancer cells. The reactions catalyzed by PGI, aldolase, GAPDH+PGK, TPI, PGAM, enolase are at near-equilibrium state, which are responsible for maintaining [F6P], [G6P], [FBP], [GA3P], [DHAP], [3-PG], [2-PG], and [PEP] in AG flux. The ΔGs for the reactions catalyzed by HK, PFK, and PK generates a major driving force for AG flux and the ΔG for the reaction catalyzed by LDH is also favorable for lactate generation by cancer cells. To investigate the role of TSAG in AG flux responding to perturbation, we used PKM2 knockdown model to investigate the interrelationship between variables (TSAG, AG rate, PK_AA_, [PKM2], [PEP], [FBP]) in the AG flux. We showed that TSAG controls [FBP] and tunes the reciprocal relationship between [PKM2] and [PEP], which stabilizes PK_AA_ in AG flux, hence AG rate is resistant to PKM2 perturbation. Thus, we reveal an important role of TSAG in maintaining the stability of AG flux and interpret the underlying biochemical mechanism. That TSAG counteracts the effect of PKM2 perturbation on AG flux could be illustrated by a schematic diagram (Figure 11). It is worth noting that the role of TSAG is to maintain intermediate concentration in the flux and to stabilize AG flux by counteracting the perturbation on AG flux.

**Figure 11.**
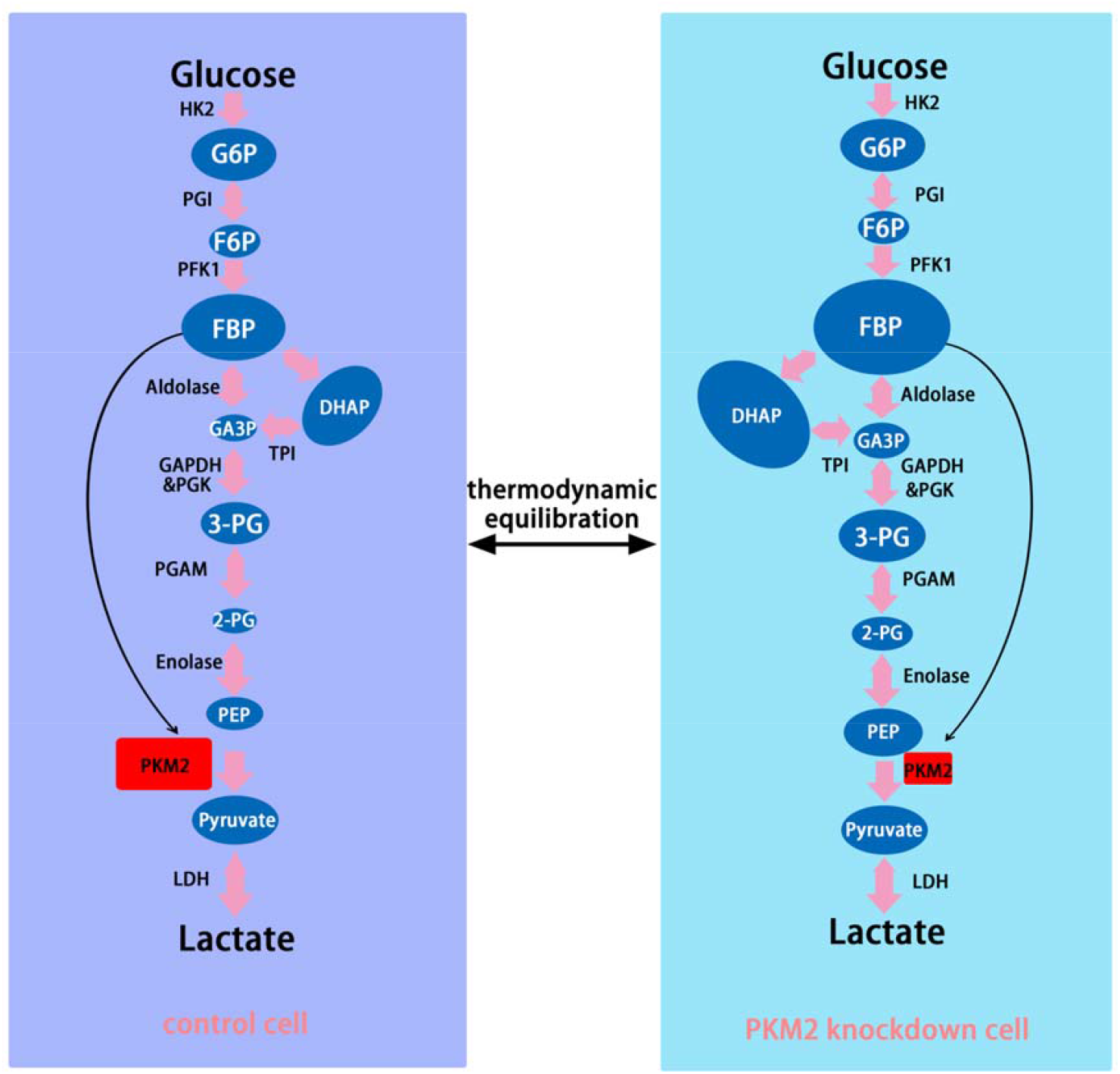
Schematic diagram to demonstrate the interaction between PKM2, AG rate, intermediate concentrations, and TSAG. Conceivably, PKM2 knockdown would cause a temporary perturbation of glycolytic flux. In response to this perturbation, the system property of the glycolytic flux – TSAG rapidly equilibrates glycolytic intermediates, characterized by the proportional increase of [PEP], [3-PG], [2-PG], [GA3P], [DHAP], and [FBP], but not [G6P], [F6P], and [pyruvate], and the unchanged ΔGs of the reactions along the pathway. Consequently, PKM2 knockdown-induced glycolytic perturbation is fixed by the thermodynamic state of the glycolytic flux. [FBP] generated from the flux saturates the allosteric regulation of PKM2 in both control and PKM2 knockdown cells. In control cells, [PKM2] is higher and [PEP] is low, whereas in PKM2 knockdown cells, [PKM2] is low and [PEP] is higher. The reciprocal relationship between [PEP] and [PKM2] is precisely tuned by the characteristic TSAG, making equal PKM2 rates as well as the overall glycolytic rates. The sizes of the blue oval 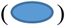 represent the relative concentrations of intermediates according to the Q values of the reactions along the glycolytic pathway; the size of the rectangle 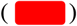 represents the relative [PKM2] in control and PKM2 knockdown cells; the pink allow 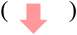 represents the rate; the arrow 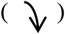 represents allosteric dominance of PKM2 by FBP.

Compared with the previous work on the regulation of AG flux by PKM2 (Anastasiou et al., 2012; Chaneton et al., 2012; Christofk, Vander Heiden, Harris, et al., 2008; Israelsen et al., 2013; Keller et al., 2014; Morgan et al., 2013; Prakasam et al., 2017), we introduce TSAG into the field and analyzes the role of TSAG in AG flux control. TSAG can effectively counteract the effect of inhibiting PKM2 on AG flux. In the PKM2 knockdown experiment, PK_SA_ decreased by 70%, but AG rate did not change significantly; in the PKM2 knockout experiment, PK_SA_ decreased by 83%, and although there was a statistically significant decrease in lactate production, the absolute amount only decreased by 13%. In other words, if there were no TSAG, PKM2 knockdown or knockout would effectively inhibit AG rate.

Earlier we proposed that thermodynamic factors play a key role in the control of AG flux (Jin et al., 2020; Zhu et al., 2021) and the entry of glucose carbon (Ying et al., 2019) into the Krebs cycle. We found that AG rate did not change significantly when PGK1 or GAPDH were treated with knockdown or inhibitors, and the underlying mechanism is that ΔG of each reaction in AG plays a critical role. The reactions catalyzed by PGK1 and GAPDH in AG are different from that catalyzed by PKM2. The reactions catalyzed by the former two in AG are in a near-equilibrium state, while the reaction catalyzed by PKM2 is far from equilibrium state, and the activity of PKM2 is regulated by many allosteric effectors in cells. After detailed reasoning and experimental procedures, we verified that thermodynamic factors also play a key role in controlling rate of PKM2 in AG flux. In line with our previous studies (Jin et al., 2020; Xie et al., 2016; Ying et al., 2019; Zhu et al., 2021), here we propose a concept that TSAG plays a key role in stabilizing AG flux.

This study has potential clinical implications. Targeting AG is a strategy for the treatment of cancer. However, the clinical translation of this strategy is not ideal at present. One of the main reasons is the unsatisfactory drug efficacy. The suboptimal efficacy is likely that the antagonism of TSAG on drug induced-perturbation of AG flux was not taken into account. Without the interference of TSAG, kinetic inhibition of glycolytic enzymes by small-molecule drugs would effectively inhibit AG flux, but TSAG can effectively counteract the kinetic inhibition of glycolytic enzymes by small-molecule drugs. Therefore, to effectively inhibit AG flux, it is necessary to simultaneously inhibit glycolytic enzymes and disrupting the antagonistic effect of TSAG on inhibiting glycolytic enzymes. Previous researches have laid a good foundation in the kinetic part of AG flux control, including the understanding of the enzymatic properties of glycolytic enzymes and the development of corresponding high-efficacious small-molecule inhibitors. On this basis, if the antagonistic effect of TSAG could be disrupted simultaneously with inhibition of glycolytic enzyme, we believe that there will be breakthroughs in the clinical translation of the treatment of cancer by targeting AG.

## Materials and methods

### Cell lines

Human lung cancer cell line A549, cervical cancer cell line HeLa, gastric cancer cell line MGC80-3, liver cancer cell line SK-HEP-1, lung cancer cell line H1299 and colon cancer cell line RKO were obtained from Cell Bank of Type Culture Collection of the Chinese Academy of Science (Shanghai, China) and were all authenticated by DNA fingerprinting. One note is that the DNA fingerprinting of MGC80-3 is not separable from that of HeLa cells. However, the morphology of HeLa and MGC80-3 were different from each other and the allostery of PK in the cell lysate from HeLa and MGC80-3 were also different from each other. Cells were cultured in RMPI-1640 medium with 10% FBS and maintained in a humidified incubator at 37°C with 5% CO_2_.

### Chemicals and enzymes

Reagents were from Sigma, including ATP (#A3377), ADP (#A5285), NAD (#N0632), NADH (#N8129), NADP (#N8035), NADPH (#N7505), Glucose (#G8270), G6P, (#G7879), F6P (#V900924), GA3P (#G5251), 3-PG (#P8877), 2-PG (#19710), PEP (#P7001), Pyruvate (#V900232), lactic acid (#L1750), HK (#H4502), PGI (#P5381), PFK (#F0137), Aldolase (#A8811), TPI (#T6258), GAPDH (#G2267), PGK (#P7634), Enolase (#E6126), PK (#P7768), LDH (#L2500), G6PDH (#G8404), α-GPDH (#G6751). FBP were purchased from aladdin (China, #F111301).

### Purification of human source recombinant PKM2

The procedure was described by us previously (J. Chen et al., 2011). Briefly, the cDNA of PKM2 was cloned into pQE-30 (Qiagen, Germany) with 6×His tag and expressed in M15 (pREP4) (AngYuBio, China). Expression was induced by 0.8 mM Isopropyl-β-D-Thiogalactoside (Beyotime, China) for 6 hours at 32 °C. The cells were collected and sonicated, and lysate was added to a Ni-NTA SefinoseTM Resin column (Sangon Biotech, China), washed with wash buffer (100 mM Tris-HCl, pH 7.6, 0.5 M NaCl and 20 mM imidazole) and eluted by elution buffer (100 mM Tris-HCl, pH 7.6, 0.5 M NaCl and 250 mM imidazole).

### PK activity assay

To determine allosteric effect of amino acid on PKM2, final concentration of 2 mM ADP, 0.2 mM PEP, 0.1 mM NADH, 5 U/ml LDH were added in 0.8 ml 37°C pre-warmed reaction buffer (200 mM HEPES, 0.5 mM EDTA, 100 mM KCl, 5 mM MgCl_2_, 5 mM Na_2_HPO_4_, pH 7.4) in a cuvette, then amino acid/FBP was added to the buffer and mixed well. Finally, cell lysate or purified recombinant PKM2/PKL was added to start the assay and absorbance at 340 nm was recorded using a spectrophotometer (DU^®^ 700, Beckman Coulter). To determine the knockdown efficiency of siRNA, PEP concentration used was 2 mM.

### Amino acid measurement using UPLC

For determination of isotopic serine and glycine, we used the same method reported by us previously (Jin et al., 2020). After 48-hour transfection, cells were cultured with glucose-free RMPI 1640 medium supplemented with 10% FBS and 6 mM [^13^C_6_]glucose for 3, 6 and 9 hours. Medium at 0, 3, 6, 9 hours were collected for LC-MS/MS measurement. Meanwhile, cells at 0, 3, 6, 9 hours were washed with PBS three times, and intracellular amino acid were extracted by adding 80% pre-cold methanol. Cells were collected by a scraper and 20,000 g centrifuge at 4°C was performed and the supernatant was evaporated by a vacuum centrifugal concentrator and was dissolved in 100 μl water. For determination of regular serine, phenylalanine, alanine, valine, proline and tryptophan in cell, cells were incubated with or without animo acid mixture (AIM) for 5 hours, and intracellular metabolites were collected as above. 10 μl dissolved metabolites or collected medium or standard was mixed with 20 μl AQC (#ab145409, abcam, amino acid derivatization agent) and 120 μl borate buffer, incubate in 60 °C for 20 min and the AQC derivatized sample was used for LC-MS/MS by a Waters Acquity UPLC system coupled to Qtrap 4000 mass spectrometer with ESI probe (Applied Biosystems Inc., Foster City, CA, USA). An AccQ-Tag Ultra RP Column was used to perform the Liquid Chromatography. The mobile phase A was 25 mM ammonia formate with 1% acetonitrile, pH 3.05 and mobile phase B was 100% acetonitrile. The gradient program was as follows: 0-0.54 min, 99.9% A, 0.1% B; 0.54-5.74 min, 99.9% A-90.9% A; 5.74-7.74 min, 90.9% A-78.8% A; 7.74-8.04 min, 78.8% A-40.4% A; 8.04-8.64 min, 40.4% A. 8.64-8.73 min, 40.4% A-99.9% A; 8.73-9.5 min, 99.9% A. 1 μl sample or standard solution was injected to perform the analysis with a flow rate at 0.7 ml/min. During the performance, the column was kept at 55°C. The following parameters were optimized and used for MASS analysis: 40 psi curtain gas, medium collision gas, 5500V ionspray voltage, temperature of the ion source 500°C, 40 psi ion source gas1 and 40 psi ion source gas2.

### Isotopic lactate determination by LC-MS/MS

Isotopic lactate tracing is based on our previously reported method(Ying et al., 2019). [^13^C_6_]Glucose was purchased from Sigma. 48 hours after transfection, HeLa-NC, HeLa-siPKM2 were washed with PBS twice, and cultured in glucose**-**free RPMI-1640 supplemented with 10% ultrafiltrated FBS and 6 mM [^13^C_6_]glucose for 6 hours. Then culture medium was collected and diluted 40 times with 100% acetonitrile and centrifuged at 25,000g for 10min at 4°C. Supernatant was collected for LS-MS/MS analysis according to methods reported previously by us (Ying et al., 2019; W. Zhang, Guo, Jiang, Ying, & Hu, 2017). Briefly, an ACQUITY BEH Amide column was used to perform liquid chromatography, kept at 50°C during analysis and the injection volume was 7.5 μl. Mobile phase A was 10 mM ammonium acetate **i**n 85% acetonitrile, 15% water, pH 9.0, and mobile phase B was 10 mM ammonium acetate in 50% acetonitrile, 50% water, pH 9.0. The gradient program was as follows: 0–0.4 min, 100% A; 0.4–2 min, 100–30% A; 2–2.5 min, 30–15% A; 2.5–3 min, 15% A; 3–3.1 min, 15–100% A; 3.1**-**7.5 min, 100% A. A 4000 QTRAP mass spectrometer (AB SCIEX) equipped with an ESI ion source (Turbospray) operated in negative ion mode was used for MS detection and same parameter setting were employed(Ying et al., 2019).

### Determination of NAD, NADH, ADP & ATP in cell

Cells in 6-well plates were washed with ice-cold PBS twice, and 0.6 ml 80% (vol/vol) pre-cold (−20°C) methanol was added per well to extract the intracellular metabolites. Then a scraper was used to collect the cells and the cell debris was discarded by 20,000 g centrifuge at 4°C. The supernatant was evaporated by a Vacuum centrifugal concentrator and was dissolved in 100μl water for following UPLC analysis. Waters ACQUITY UPLC system with an ACQUITY UPLC HSS T3 column was used to perform the Liquid Chromatography. The mobile phase A was 20 mM Triethylamine in 99%/1% water/acetonitrile (pH 6.5) and mobile phase B was 100% acetonitrile. The gradient program was as follows: 0-9 min, 100% A-90% A; 9-10 min, 90% A-0% A; 10-11 min, 100% B; 11-12 min, 0% A-100% A. 12-20 min, 100% A.10 μl sample or standard solution was injected to perform the analysis with a flow rate at 0.3 ml/min. During the performance, the column was kept at 40°C.

### In vitro Cell-free system model for glycolysis

Previously we had described an in vitro cell-free system as a glycolysis model (Xie et al., 2016; Xie et al., 2014). We used the reaction buffer containing 200 mM HEPES, 0.5 mM EDTA, 100 mM KCl, 5 mM MgCl_2_, 5 mM Na_2_HPO_4_, 4 mM ADP, 1.5 mM ATP, 5 mM glucose, 0.1 mM NADH and 2 mM NAD for this glycolysis system. 70 μl lysate (8-10 μg/μl protein) was added to 630 μl reaction buffer, and the mixture was incubated at 37 °C for 30 minutes. Then we added 600 μl 1 M HClO_4_ to the mixture to terminate the reaction and later 100 μl 3 M K_2_CO_3_ was added to neutralize the buffer. The mixture was kept on ice for 20 min further. Supernatant was obtained by 10,000 g centrifuge at 4 °C and was used for glycolytic intermediates determination.

### Determination of glycolytic intermediates, glucose and lactate

The same enzymatic methods to determine glucose, lactate and glycolytic intermediates were described by us previously (Jin et al., 2020; Xie et al., 2016; Zhu et al., 2021).

### Determination of glycolytic enzyme activity

We determined glycolytic enzyme activity at saturating substrate concentration according to previously reported methods (Passonneau, 1993) with some modifications (Jin et al., 2020; Xie et al., 2016; Zhu et al., 2021). Briefly, 0.8 ml reaction buffer was added to the cuvette, and substrates were added. The reaction was started by adding cell lysate and mixed, and then absorbance at 340 nm was recorded using a spectrophotometer (DU^®^ 700, Beckman Coulter).

### Determination of intracellular glycolytic intermediates by cycling method

Negative control or siPKM2 transfected cells were washed by ice cold PBS twice, and 600 μl 1 M pre-cold HClO_4_ was added to every 5 wells of 6-well plate. Cells were collected by a scraper, incubated on ice for 30 min, and neutralized by 100 μl 3 M K_2_CO_3_. Then supernatant was obtained by 10,000 g centrifuge at 4°C. Then 50 μl 2 M NaOH was added to the supernatant and kept at 60°C for 5 minutes, and neutralized again by adding 50 μl 2 M HCl. These two steps eliminated intracellular NAD(H) / NADP(H) possibly. Meanwhile, a same sixth well of cells was trypsinized and collected to determine the cell number and cell size by a cell counter (JIMBIO). After these sample preparing procedures, intracellular glycolytic intermediates were determined by cycling method as previously described by us (Jin et al., 2020; Zhu et al., 2021).

### siRNA transfection to knockdown PKM2

1.6×10^5^ cells were seeded into each well of 6-well plates and cultured overnight. Cells were transfected using Lipofectamine 3000 (Thermo Fisher Scientific) according to manufacturer’s protocol, with either negative control siRNA (NC) or siPKM2 (Ribobio, China). The siRNA sequences were as follows: siPKM2, sense, GUGGUGAUCUAGGCAUUGAdTdT; antisense, UCAAUGCCUAGAUCACCAC dTdT; NC, sense, UUCUCCGAACGUGUCACGUdTdT; antisense, ACGUGACACGUUCGGAGAA dTdT. 48 hours after transfection, cells were washed with PBS and 2 ml fresh complete RMPI-1640 plus 8 mM glucose were added to each well. Then we collected 10 μL media at 1, 2, 3, 4, 5, 6 hours and determined glucose & lactate afterwards using the method described by us previously (Jin et al., 2020). The cells were counted and collected for enzyme activity assay, western blot, or intracellular intermediates determination.

### Knockout of PKM2 using Crispr/Cas9 system

PKM2 was knocked out using Crispr/Cas 9 system according to protocol in Feng Zhang’s lab (Ran et al., 2013). Briefly, PKM2 sgRNA (Forward: CACCGCATTCTTGATGGTCTCCGCA; Reverse: AAACTGCGGAGACCATCAAGAATGC) was designed and annealed, ligated to BbsI digested backbone plasmid (Addgene, #62988), and transfected into cells using Lipofectamine 3000 (Invitrogen, #L3000015). HeLa cells transfected with expression plasmid without sgRNA insertion were used as a control. Then cells were selected with 2 μg/ml puromycin for 48 hours, allowed to grow to 80% confluence and plated to 96 well plates in order to obtain clones from single cells. The clones were validated using western blot and enzyme activity assay.

### Cell cycle

Cell cycle assay was performed using a Cell cycle staining Kit (# 70-CCS012, MultiSciences, China) according to manufacturer’s protocol. Briefly, HeLa cells were transfected with NC or siPKM2 for 48 hours, trypsinized and seeded to a new 6-well plate overnight, then collected and subjected to flow cytometer analysis.

### Western blot

Cells were washed with cold PBS, then lysed with M-PER™ Mammalian Protein Extraction Reagent (Thermo Fisher Scientific) supplemented with cocktail (MedChemExpress) on ice for 30 minutes. Protein concentration was determined using BCA protein assay kit (Thermo Fisher Scientific). The protein was boiled for 5 minutes with loading buffer and 20 μg was subjected to 10% SDS-PAGE, transferred to PVDF membrane, and incubated with primary body PKM2 (HuaBio, #ER1802-70). GAPDH (Proteintech, #60004-1-lg) was used as internal control.

### Calculation of the Gibbs free energy change ΔG of glycolytic reactions

ΔG was calculated as previously reported (Jin et al., 2020) according to the equation

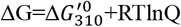

where 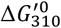 is the standard transformed Gibbs free energy at 37°C and Q was calculated according to intermediate concentrations and was listed in Supplementary Tables. NAD/NADH was set as 78 according to our previously reported study (Xie et al., 2016) and [Pi] in the cell was 1.5 mM according to (Giachelli, 2003). According to ΔG=ΔH-TΔS, since the change of ΔH and ΔS is negligible between 37°C and 25°C(Mendez, 2008), we deduced the equation to new form that

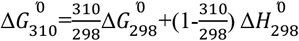

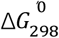 and 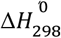 are available in references(Alberty, 2006; Keleti, Foldi, Erdei, & Tro, 1972; Li, Wu, Qi, & Beard, 2011; Varga et al., 2009).

### Statistical analysis

All experiments were repeated at least 3 times, and all data were analyzed using GraphPad Prism7. For comparisons of two groups, two-tailed Student’s t test was performed.

### Data availability

All data are in the article.

## Abbreviatoions

3-PG: 3-phosphoglycerate
2-PG: 2-phosphoglycerate
AIM: allosteric inhibitor mixture
AG: Aerobic glycolysis
Ala: Alanine
F6P: fructose 6-phosphate
FBP: fructose 1,6-bisphosphate
FBS: fetal bovine serum
G6P: glucose 6-phosphate
G6PDH: glucose-6-phosphate dehydrogenase
GA3P: glyceraldehyde 3-phosphate
GAPDH: glyceraldehyde 3-phosphate dehydrogenase
ΔG: Gibbs free energy
Glc: glucose
HK: hexokinase
LDH: lactate dehydrogenase
MTS: methanethiosulfonate
NC: negative control
PEP: phosphoenolpyruvate
PES: phenazine ethosulfate
PFK: phosphofructokinase
PGI: phosphohexose isomerase
PGK: phosphoglycerate kinase
Phe: phenylalanine
PK: pyruvate kinase
Pro: proline
Pyr: pyruvate
Q: reaction quotient
Ser: serine
TPI: triose phosphate isomerase
Trp: tryptophan
[x], the concentration of x: e.g., [PK] refers the concentration of PK
TSAG: thermodynamic state of AG
UPLC: ultra performance liquid chromatography
Val: valine

## Author contributions

X.H. conceived the conception, designed the study, interpreted the data, and wrote the paper. C.J., W.H., Y.W., H. W., S.Z., and M.Y. conducted the experiments.

## Funding and additional information

This work has been supported in part China Natural Science Foundation projects (82073038, 81772947), a key project (2018C03009) funded by Zhejiang Provincial Department of Sciences and Technologies, the Fundamental Research Funds for the Central Universities (2017XZZX001-01, 2019FZJD009), the National Ministry of Education, China, to X.H.

## Conflict of interest

The authors declare that they have no conflicts of interest with the contents of this article.

## Supplementary information

### Supplementary Table

**Supplementary Table 1.** Glycolytic intermediates (mM) in 6 cancer cells. Date represent the mean ± SD of 3 independent experiments performed in triplicates.

|  | Glc | G6P | F6P | FBP | DHAP | GA3P | 3-PG | 2-PG | PEP | Pyruvate |
| --- | --- | --- | --- | --- | --- | --- | --- | --- | --- | --- |
| A549 | 2.2±0.1 | 0.14±0.03 | 0.10±0.01 | 0.39±0.02 | 0.36±0.04 | 0.06±0.02 | 0.12±0.02 | 0.04±0.01 | 0.04±0.01 | 0.13±0.04 |
| H1299 | 4.3±0.2 | 0.17±0.02 | 0.05±0.01 | 0.22±0.04 | 0.34±0.11 | 0.04±0.00 | 0.24±0.02 | 0.04±0.01 | 0.04±0.01 | 0.22±0.03 |
| HeLa | 3.7±0.5 | 0.43±0.02 | 0.16±0.03 | 0.63±0.10 | 0.54±0.09 | 0.06±0.01 | 0.30±0.08 | 0.09±0.05 | 0.07±0.03 | 0.44±0.06 |
| MGC80-3 | 1.5±0.0 | 0.08±0.02 | 0.04±0.00 | 0.36±0.02 | 0.27±0.01 | 0.04±0.00 | 0.23±0.02 | 0.08±0.03 | 0.14±0.01 | 0.17±0.02 |
| RKO | 2.6±0.1 | 0.22±0.02 | 0.13±0.00 | 0.21±0.02 | 0.21±0.03 | 0.04±0.01 | 0.20±0.03 | 0.06±0.01 | 0.05±0.02 | 0.23±0.07 |
| SK-HEP-1 | 2.6±0.1 | 0.25±0.02 | 0.13±0.02 | 0.80±0.11 | 0.44±0.14 | 0.06±0.01 | 0.34±0.06 | 0.13±0.04 | 0.09±0.01 | 0.33±0.04 |

**Supplementary Table 2.**
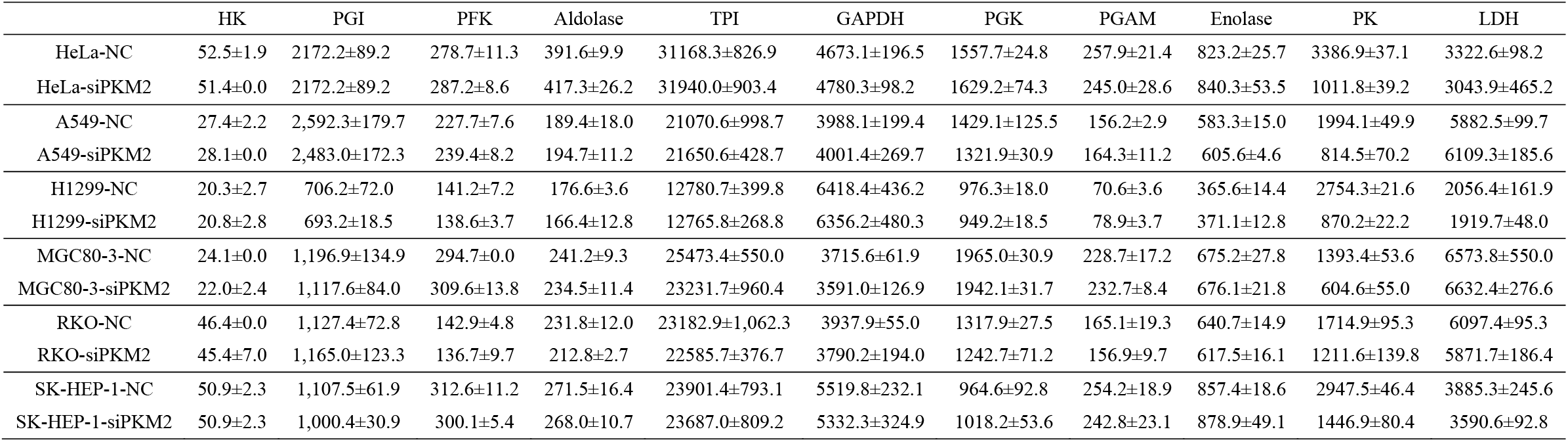
Specific activities (nmol/min.mg protein) of glycolytic enzymes with or without PKM2 knockdown in 6 cancer cell lines. Date represent the mean ± SD of 3 independent experiments performed in triplicates.

|  | HK | PGI | PFK | Aldolase | TPI | GAPDH | PGK | PGAM | Enolase | PK | LDH |
| --- | --- | --- | --- | --- | --- | --- | --- | --- | --- | --- | --- |
| HeLa-NC | 52.5±1.9 | 2172.2±89.2 | 278.7±11.3 | 391.6±9.9 | 31168.3±826.9 | 4673.1±196.5 | 1557.7±24.8 | 257.9±21.4 | 823.2±25.7 | 3386.9±37.1 | 3322.6±98.2 |
| HeLa-siPKM2 | 51.4±0.0 | 2172.2±89.2 | 287.2±8.6 | 417.3±26.2 | 31940.0±903.4 | 4780.3±98.2 | 1629.2±74.3 | 245.0±28.6 | 840.3±53.5 | 1011.8±39.2 | 3043.9±465.2 |
| A549-NC | 27.4±2.2 | 2,592.3±179.7 | 227.7±7.6 | 189.4±18.0 | 21070.6±998.7 | 3988.1±199.4 | 1429.1±125.5 | 156.2±2.9 | 583.3±15.0 | 1994.1±49.9 | 5882.5±99.7 |
| A549-siPKM2 | 28.1±0.0 | 2,483.0±172.3 | 239.4±8.2 | 194.7±11.2 | 21650.6±428.7 | 4001.4±269.7 | 1321.9±30.9 | 164.3±11.2 | 605.6±4.6 | 814.5±70.2 | 6109.3±185.6 |
| H1299-NC | 20.3±2.7 | 706.2±72.0 | 141.2±7.2 | 176.6±3.6 | 12780.7±399.8 | 6418.4±436.2 | 976.3±18.0 | 70.6±3.6 | 365.6±14.4 | 2754.3±21.6 | 2056.4±161.9 |
| H1299-siPKM2 | 20.8±2.8 | 693.2±18.5 | 138.6±3.7 | 166.4±12.8 | 12765.8±268.8 | 6356.2±480.3 | 949.2±18.5 | 78.9±3.7 | 371.1±12.8 | 870.2±22.2 | 1919.7±48.0 |
| MGC80-3-NC | 24.1±0.0 | 1,196.9±134.9 | 294.7±0.0 | 241.2±9.3 | 25473.4±550.0 | 3715.6±61.9 | 1965.0±30.9 | 228.7±17.2 | 675.2±27.8 | 1393.4±53.6 | 6573.8±550.0 |
| MGC80-3-siPKM2 | 22.0±2.4 | 1,117.6±84.0 | 309.6±13.8 | 234.5±11.4 | 23231.7±960.4 | 3591.0±126.9 | 1942.1±31.7 | 232.7±8.4 | 676.1±21.8 | 604.6±55.0 | 6632.4±276.6 |
| RKO-NC | 46.4±0.0 | 1,127.4±72.8 | 142.9±4.8 | 231.8±12.0 | 23182.9±1,062.3 | 3937.9±55.0 | 1317.9±27.5 | 165.1±19.3 | 640.7±14.9 | 1714.9±95.3 | 6097.4±95.3 |
| RKO-siPKM2 | 45.4±7.0 | 1,165.0±123.3 | 136.7±9.7 | 212.8±2.7 | 22585.7±376.7 | 3790.2±194.0 | 1242.7±71.2 | 156.9±9.7 | 617.5±16.1 | 1211.6±139.8 | 5871.7±186.4 |
| SK-HEP-1-NC | 50.9±2.3 | 1,107.5±61.9 | 312.6±11.2 | 271.5±16.4 | 23901.4±793.1 | 5519.8±232.1 | 964.6±92.8 | 254.2±18.9 | 857.4±18.6 | 2947.5±46.4 | 3885.3±245.6 |
| SK-HEP-1-siPKM2 | 50.9±2.3 | 1,000.4±30.9 | 300.1±5.4 | 268.0±10.7 | 23687.0±809.2 | 5332.3±324.9 | 1018.2±53.6 | 242.8±23.1 | 878.9±49.1 | 1446.9±80.4 | 3590.6±92.8 |

**Supplementary Table 3.**
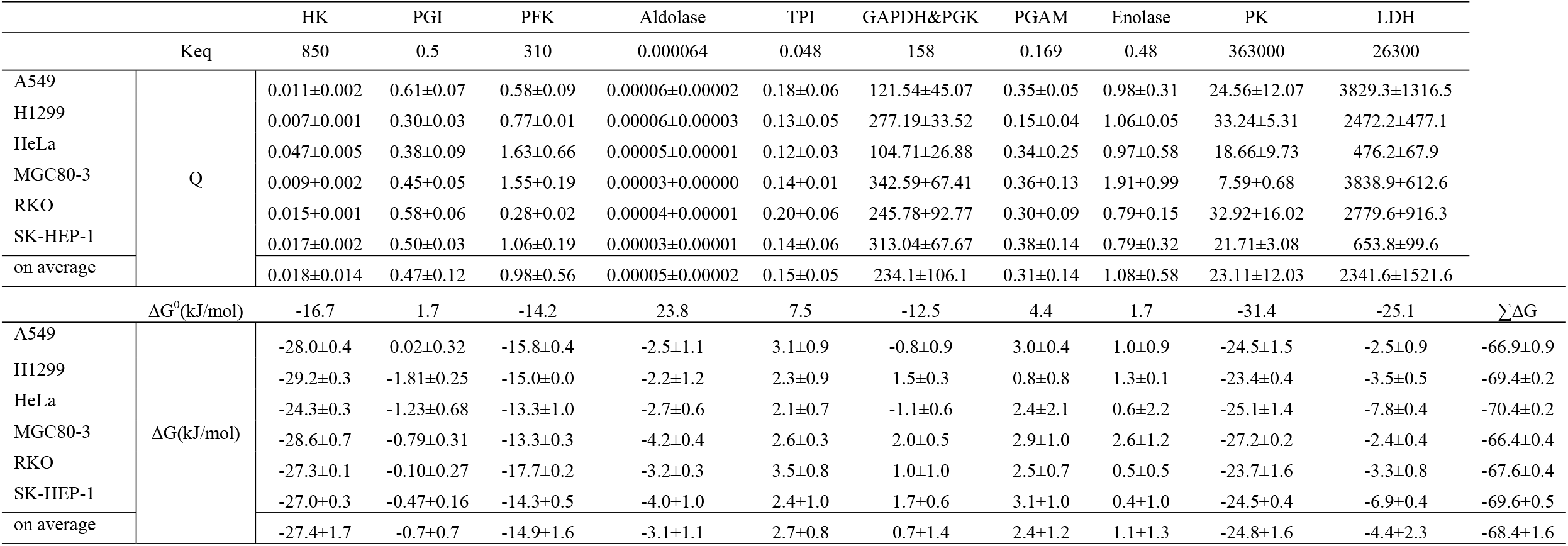
Q and ΔG of glycolytic enzymes in cancer cells. Qs were calculated according to the data in Supplementary Table 1. ΔG values in the lower row (HeLa-NC&HeLa-siPKM2) were calculated according to the data in Supplementary table 5. Data represent the mean ± SD of 3 independent experiments.

|  |  | HK | PGI | PFK | Aldolase | TPI | GAPDH&PGK | PGAM | Enolase | PK | LDH |  |
| --- | --- | --- | --- | --- | --- | --- | --- | --- | --- | --- | --- | --- |
| K<sub>eq</sub> |  | 850 | 0.5 | 310 | 0.000064 | 0.048 | 158 | 0.169 | 0.48 | 363000 | 26300 |  |
| A549 | Q | 0.011±0.002 | 0.61±0.07 | 0.58±0.09 | 0.00006±0.00002 | 0.18±0.06 | 121.54±45.07 | 0.35±0.05 | 0.98±0.31 | 24.56±12.07 | 3829.3±1316.5 | |
| H1299 | | 0.007±0.001 | 0.30±0.03 | 0.77±0.01 | 0.00006±0.00003 | 0.13±0.05 | 277.19±33.52 | 0.15±0.04 | 1.06±0.05 | 33.24±5.31 | 2472.2±477.1 | |
| HeLa | | 0.047±0.005 | 0.38±0.09 | 1.63±0.66 | 0.00005±0.00001 | 0.12±0.03 | 104.71±26.88 | 0.34±0.25 | 0.97±0.58 | 18.66±9.73 | 476.2±67.9 | |
| MGC80-3 | | 0.009±0.002 | 0.45±0.05 | 1.55±0.19 | 0.00003±0.00000 | 0.14±0.01 | 342.59±67.41 | 0.36±0.13 | 1.91±0.99 | 7.59±0.68 | 3838.9±612.6 | |
| RKO | | 0.015±0.001 | 0.58±0.06 | 0.28±0.02 | 0.00004±0.00001 | 0.20±0.06 | 245.78±92.77 | 0.30±0.09 | 0.79±0.15 | 32.92±16.02 | 2779.6±916.3 | |
| SK-HEP-1 | | 0.017±0.002 | 0.50±0.03 | 1.06±0.19 | 0.00003±0.00001 | 0.14±0.06 | 313.04±67.67 | 0.38±0.14 | 0.79±0.32 | 21.71±3.08 | 653.8±99.6 | |
| on average | | 0.018±0.014 | 0.47±0.12 | 0.98±0.56 | 0.00005±0.00002 | 0.15±0.05 | 234.1±106.1 | 0.31±0.14 | 1.08±0.58 | 23.11±12.03 | 2341.6±1521.6 | |
| $\Delta G^0$(kJ/mol) | | -16.7 | 1.7 | -14.2 | 23.8 | 7.5 | -12.5 | 4.4 | 1.7 | -31.4 | -25.1 | $\Sigma \Delta G$ |
| A549 | $\Delta G$(kJ/mol) | -28.0±0.4 | 0.02±0.32 | -15.8±0.4 | -2.5±1.1 | 3.1±0.9 | -0.8±0.9 | 3.0±0.4 | 1.0±0.9 | -24.5±1.5 | -2.5±0.9 | -66.9±0.9 |
| H1299 | | -29.2±0.3 | -1.81±0.25 | -15.0±0.0 | -2.2±1.2 | 2.3±0.9 | 1.5±0.3 | 0.8±0.8 | 1.3±0.1 | -23.4±0.4 | -3.5±0.5 | -69.4±0.2 |
| HeLa | | -24.3±0.3 | -1.23±0.68 | -13.3±1.0 | -2.7±0.6 | 2.1±0.7 | -1.1±0.6 | 2.4±2.1 | 0.6±2.2 | -25.1±1.4 | -7.8±0.4 | -70.4±0.2 |
| MGC80-3 | | -28.6±0.7 | -0.79±0.31 | -13.3±0.3 | -4.2±0.4 | 2.6±0.3 | 2.0±0.5 | 2.9±1.0 | 2.6±1.2 | -27.2±0.2 | -2.4±0.4 | -66.4±0.4 |
| RKO | | -27.3±0.1 | -0.10±0.27 | -17.7±0.2 | -3.2±0.3 | 3.5±0.8 | 1.0±1.0 | 2.5±0.7 | 0.5±0.5 | -23.7±1.6 | -3.3±0.8 | -67.6±0.4 |
| SK-HEP-1 | | -27.0±0.3 | -0.47±0.16 | -14.3±0.5 | -4.0±1.0 | 2.4±1.0 | 1.7±0.6 | 3.1±1.0 | 0.4±1.0 | -24.5±0.4 | -6.9±0.4 | -69.6±0.5 |
| on average | | -27.4±1.7 | -0.7±0.7 | -14.9±1.6 | -3.1±1.1 | 2.7±0.8 | 0.7±1.4 | 2.4±1.2 | 1.1±1.3 | -24.8±1.6 | -4.4±2.3 | -68.4±1.6 |

**Supplementary Table 4.** Intracellular glycolytic intermediates in cancer cells with or without PKM2 knockdown. Data are mean ± SD, n=3. Results were confirmed by two independent experiments.

|  | Glc | G6P | F6P | FBP | DHAP | GA3P | 3-PG | 2-PG | PEP | Pyr | NAD | NADH | ADP | ATP |
| --- | --- | --- | --- | --- | --- | --- | --- | --- | --- | --- | --- | --- | --- | --- |
| A549-NC | 2.2±0.1 | 0.14±0.03 | 0.10±0.01 | 0.39±0.02 | 0.36±0.04 | 0.06±0.02 | 0.12±0.02 | 0.04±0.01 | 0.04±0.01 | 0.13±0.04 | 1.5±0.1 | 0.11±0.04 | 0.5±0.1 | 3.5±0.2 |
| A549-siPKM2 | 2.1±0.1 | 0.12±0.01 | 0.08±0.00 | 0.55±0.01 | 0.49±0.03 | 0.08±0.01 | 0.18±0.00 | 0.06±0.01 | 0.07±0.01 | 0.11±0.02 | 1.8±0.2 | 0.19±0.04 | 0.7±0.1 | 4.3±0.9 |
| H1299-NC | 4.3±0.2 | 0.17±0.02 | 0.05±0.01 | 0.22±0.04 | 0.34±0.11 | 0.04±0.00 | 0.24±0.02 | 0.04±0.01 | 0.04±0.01 | 0.22±0.03 | 1.6±0.1 | 0.05±0.01 | 0.9±0.1 | 5.0±0.3 |
| H1299-siPKM2 | 4.4±0.1 | 0.19±0.02 | 0.07±0.03 | 0.64±0.03 | 0.60±0.05 | 0.07±0.01 | 0.31±0.04 | 0.07±0.01 | 0.07±0.01 | 0.23±0.01 | 1.8±0.2 | 0.06±0.02 | 1.1±0.1 | 6.3±0.9 |
| HeLa-NC | 3.7±0.5 | 0.43±0.02 | 0.16±0.03 | 0.63±0.10 | 0.54±0.09 | 0.06±0.01 | 0.30±0.08 | 0.09±0.05 | 0.07±0.03 | 0.44±0.06 | 4.0±0.2 | 0.66±0.02 | 2.0±0.1 | 5.1±0.3 |
| HeLa-siPKM2 | 2.9±0.2 | 0.36±0.05 | 0.11±0.01 | 1.01±0.21 | 0.95±0.15 | 0.09±0.02 | 0.50±0.09 | 0.19±0.03 | 0.21±0.02 | 0.44±0.04 | 3.8±0.1 | 0.66±0.06 | 2.2±0.1 | 4.8±0.2 |
| MGC80-3-NC | 1.5±0.0 | 0.08±0.02 | 0.04±0.00 | 0.36±0.02 | 0.27±0.01 | 0.04±0.00 | 0.23±0.02 | 0.08±0.03 | 0.14±0.01 | 0.17±0.02 | 1.1±0.0 | 0.03±0.01 | 0.7±0.0 | 4.3±0.2 |
| MGC80-2-siPKM2 | 1.4±0.0 | 0.09±0.01 | 0.04±0.00 | 0.51±0.01 | 0.38±0.02 | 0.06±0.01 | 0.39±0.03 | 0.07±0.02 | 0.23±0.02 | 0.16±0.05 | 1.0±0.0 | 0.04±0.04 | 0.6±0.1 | 4.2±0.2 |
| RKO-NC | 2.6±0.1 | 0.22±0.02 | 0.13±0.00 | 0.21±0.02 | 0.21±0.03 | 0.04±0.01 | 0.20±0.03 | 0.06±0.01 | 0.05±0.02 | 0.23±0.07 | 1.5±0.0 | 0.09±0.01 | 1.0±0.1 | 6.1±0.2 |
| RKO-siPKM2 | 2.5±0.0 | 0.24±0.01 | 0.11±0.02 | 0.28±0.01 | 0.33±0.01 | 0.07±0.01 | 0.31±0.01 | 0.07±0.01 | 0.11±0.02 | 0.19±0.03 | 1.6±0.1 | 0.06±0.04 | 1.2±0.2 | 6.6±0.5 |
| SK-HEP-1-NC | 2.6±0.1 | 0.25±0.02 | 0.13±0.02 | 0.80±0.11 | 0.44±0.14 | 0.06±0.01 | 0.34±0.06 | 0.13±0.04 | 0.09±0.01 | 0.33±0.04 | 1.6±0.1 | 0.22±0.11 | 1.4±0.1 | 8.3±0.5 |
| SK-HEP-1-siPKM2 | 2.7±0.1 | 0.24±0.02 | 0.10±0.04 | 1.51±0.28 | 1.44±0.21 | 0.12±0.02 | 1.07±0.19 | 0.14±0.02 | 0.15±0.03 | 0.31±0.08 | 1.6±0.1 | 0.11±0.06 | 1.5±0.1 | 8.3±0.5 |

**Supplementary Table 5.**
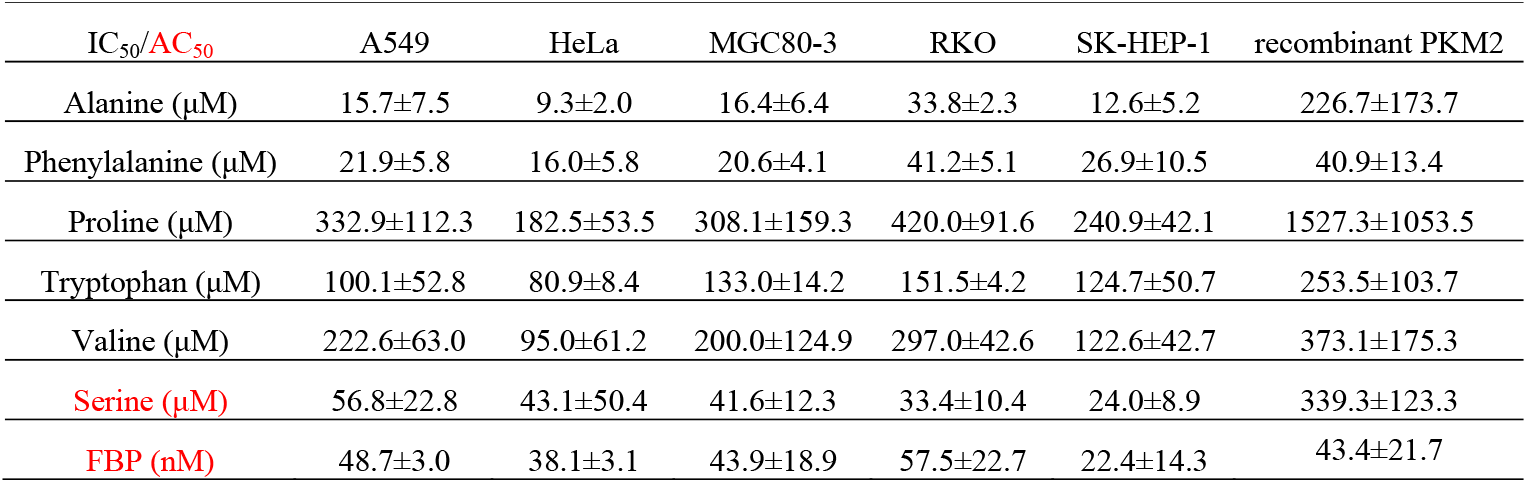
IC_50_ of allosteric inhibitor and AC_50_ of FBP and serine for cell lysate PK and pure recombinant PKM2. Data were estimated according to the kinetic curves of Supplementary Figure 1 and represent the mean ± SD of 3 independent experiments performed in triplicates.

| IC<sub>50</sub>/AC<sub>50</sub> | A549 | HeLa | MGC80-3 | RKO | SK-HEP-1 | recombinant PKM2 |
| --- | --- | --- | --- | --- | --- | --- |
| Alanine (μM) | 15.7±7.5 | 9.3±2.0 | 16.4±6.4 | 33.8±2.3 | 12.6±5.2 | 226.7±173.7 |
| Phenylalanine (μM) | 21.9±5.8 | 16.0±5.8 | 20.6±4.1 | 41.2±5.1 | 26.9±10.5 | 40.9±13.4 |
| Proline (μM) | 332.9±112.3 | 182.5±53.5 | 308.1±159.3 | 420.0±91.6 | 240.9±42.1 | 1527.3±1053.5 |
| Tryptophan (μM) | 100.1±52.8 | 80.9±8.4 | 133.0±14.2 | 151.5±4.2 | 124.7±50.7 | 253.5±103.7 |
| Valine (μM) | 222.6±63.0 | 95.0±61.2 | 200.0±124.9 | 297.0±42.6 | 122.6±42.7 | 373.1±175.3 |
| Serine (μM) | 56.8±22.8 | 43.1±50.4 | 41.6±12.3 | 33.4±10.4 | 24.0±8.9 | 339.3±123.3 |
| FBP (nM) | 48.7±3.0 | 38.1±3.1 | 43.9±18.9 | 57.5±22.7 | 22.4±14.3 | 43.4±21.7 |

**Supplementary Table 6.** Glycolytic intermediates (mM) in cell-free system. Data are mean ± SD, n=3. Results were confirmed by two independent experiments.

|  | Glc | G6P | F6P | FBP | DHAP | GA3P | 3-PG | 2-PG | PEP | Pyr | Lactate |
| --- | --- | --- | --- | --- | --- | --- | --- | --- | --- | --- | --- |
| HeLa-NC | 4.20±0.09 | 0.021±0.00 | 0.007±0.000 | 0.034±0.00 | 0.094±0.00 | 0.003±0.001 | 0.023±0.003 | 0.004±0.000 | 0.010±0.001 | 0.008±0.001 | 1.42±0.02 |
| HeLa-siPKM2 | 4.16±0.10 | 0.022±0.00 | 0.006±0.001 | 0.039±0.00 | 0.108±0.00 | 0.008±0.002 | 0.048±0.002 | 0.006±0.001 | 0.016±0.000 | 0.007±0.000 | 1.43±0.08 |

### Supplementary Figures

**Supplementary Figure 1.**
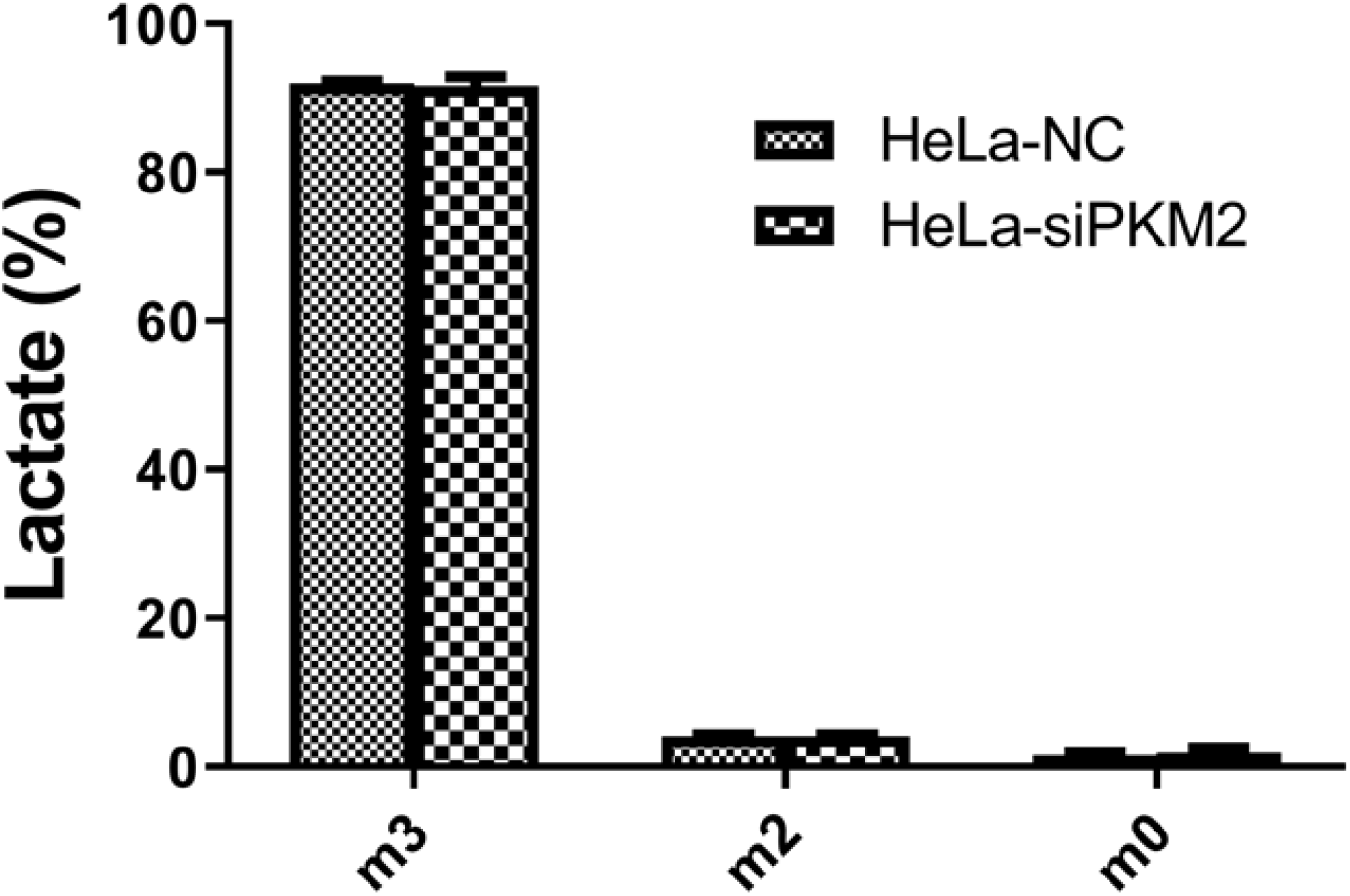
Tracing glucose carbon to lactate. Cells were incubated with 6 mM [^13^C_6_]glucose for 6 hours, and the percentage of the generated lactate isotopologues were measured by LC-MS. Data are mean ± SD, n=3. Results were confirmed by two independent experiments.

**Supplementary Figure 2.**
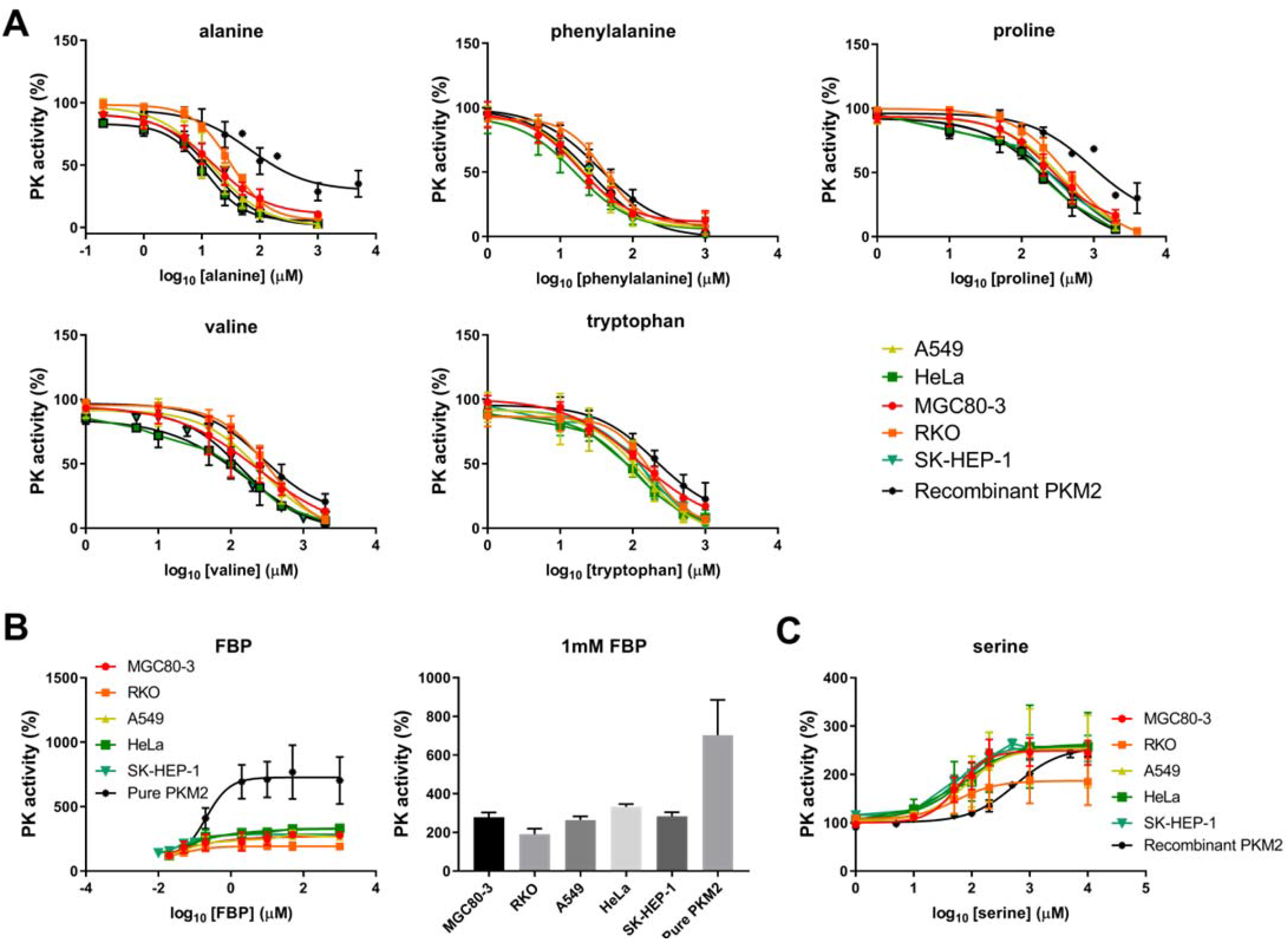
Effects of amino acids and FBP that allosterically inhibit or activate PK. **(A)** Kinetic profiles of cell lysate PK or pure recombinant PKM2 determined at different concentrations of alanine, phenylalanine, proline, tryptophan or valine. **(B)** Left panel, kinetic profiles of cell lysate PK or pure recombinant PKM2 determined at different concentrations of FBP; right panel, PK activities at 1mM FBP were extracted and presented. **(C)** kinetic profiles of cell lysate PK or recombinant PKM2 determined at different concentration of serine. Pyruvate kinase activity was assayed at 2 mM ADP and 0.2 mM PEP. PK activities with no activators were used as control (100%). Data represent the mean ± S.D. of 3 independent experiments performed in triplicates.

## Notes

### Competing Interest Statement

The authors have declared no competing interest.

